# The geometry of decision-making

**DOI:** 10.1101/2021.05.26.445795

**Authors:** Vivek Hari Sridhar, Liang Li, Dan Gorbonos, Máté Nagy, Bianca R. Schell, Timothy Sorochkin, Nir S. Gov, Iain D. Couzin

## Abstract

Choosing among spatially-distributed options is a central challenge for animals, from deciding among alternative potential food sources or refuges, to choosing with whom to associate. Using an integrated theoretical and experimental approach (employing immersive virtual reality), we consider the interplay between movement and vectorial integration during decision-making regarding two, or more, options in space. In computational models of this process we reveal the occurrence of spontaneous and abrupt “critical” transitions (associated with specific geometrical relationships) whereby organisms spontaneously switch from averaging vectorial information among, to suddenly excluding one, among the remaining options. This bifurcation process repeats until only one option—the one ultimately selected—remains. Thus we predict that the brain repeatedly breaks multi-choice decisions into a series of binary decisions in space-time. Experiments with fruit flies, desert locusts, and larval zebrafish reveal that they exhibit these same bifurcations, demonstrating that across taxa and ecological context, there exist fundamental geometric principles that are essential to explain how, and why, animals move the way they do.

---

Animals constantly face the need to make decisions, and many such decisions require choosing among multiple spatially-distributed options. Despite this, most studies have focused on the outcome of decisions (1–3) (i.e. which option among alternatives is chosen), as well as the time taken to make decisions (4–6), but seldom on the movement of animals throughout the decision-making process. Motion is, however, crucial in terms of how space is represented by organisms during spatial decision-making; the brains of a wide range of species, from insects (7, 8) to vertebrates (9, 10), have been shown to represent egocentric spatial relationships, such as the position of desired targets, via explicit vectorial representation (11, 12). Such neuronal representations must, and do, change as animals move through space. Thus, while the movement of an animal may, initially, appear to simply be a readout of the decision made by the brain—and consequently not particularly informative—this view overlooks important dynamical properties introduced into the decision-making process that result from the inevitable time-varying geometrical relationships between an organism and spatially-distributed options (i.e. potential ‘targets’ in space).

Due to a dearth of existing studies, and with the objective to develop the necessary foundational understanding of the ‘geometry’ of decision-making, we focus here—first theoretically and then experimentally—on the consequences of the recursive interplay between movement and (collective) vectorial integration in the brain during relatively simple spatial decisions. We employ immersive virtual reality to investigate decision-making regarding multiple (two or more) options in both invertebrate (the fruit fly *Drosophila melanogaster*, and desert locust *Schistocerca gregaria*) and vertebrate (larval zebrafish *Danio rerio*) models. Doing so allows us to reveal the emergence of geometric principles that transcend the study organism and the decision-making context, and thus are expected to be broadly relevant across taxa. In support of this finding we also explore how these principles extend to collective decision-making in mobile animal groups, allowing us to gain insights across three scales of biological organisation, from neural dynamics, to both individual and collective decision-making.

## Modelling decision-making on the move

Congruent with neurobiological studies of the invertebrate and vertebrate brain, we consider organisms to have an egocentric vectorial representation of spatial options (11–13). We then consider the collective dynamics of vector integration in the brain assuming there exists reinforcement (excitation/positive feedback) among neural ensembles that have similar directional representations (goal vectors), and global inhibition and/or negative feedback (both produce broadly similar results, see SI Appendix and Fig. S1) among neural ensembles that differ in vectorial representation. This captures, in a simple mathematical formulation, the essence of both explicit ring-attractor networks (as found in insects (7)), and computation among competing neural groups (as in the mammalian brain (14)). The animal’s relative preference for a target is given by activity of neurons that encode direction to that target relative to activity of neurons that encode direction to other targets, and the angular sensitivity of the neural representations (angular difference at which excitation no longer occurs) is specified by a neural tuning parameter, *ν*. The network then computes a unique ‘consensus’ vector (‘activity bump’) that, along with some angular noise, represents the animal’s desired direction of movement (Fig. S2). This is then translated into motor output (see SI Appendix for model details \cite{pinkoviezky_collective_2018}). Stochasticity in neural dynamics is implemented here as the neural noise parameter, *T*.

While capturing known, generic features of neural integration, our model is deliberately minimal. This serves multiple purposes: firstly, following principles of maximum parsimony we seek to find a simple model that can both predict and explain, the observed phenomena; secondly, we aim to reveal general principles and thus consider features that are known to be valid across organisms irrespective of inevitable difference in structural organization of the brain; thirdly, it provides a convenient means to implement neural noise, and can be mapped to the class of neural ring-attractor models widely-used in neuroscience (15–18) (see SI Appendix for details). In addition, our results are shown to be extremely robust to model assumptions, suggesting that it provides an appropriate low-level description of essential system properties.

## Deciding between two options

Beginning with the simplest case, we consider the feedback between motion and internal vectorial-computation when an animal is presented with two equally-attractive, but spatially-discrete, options. In this case the activity of neurons encoding option 1, *N*_1_ will be equal to those encoding option 2, *N*_2_ (Fig. 1A). Our model predicts that an animal moving, from a relatively distant location, towards the two targets, will spontaneously compute the average directional preference, resulting in corresponding motion in a direction oriented between the two targets. As it approaches the targets, however, upon reaching a certain angular difference between the options, the internal network undergoes a sudden transition in which it spontaneously selects one, or the other, target (Fig. 1C). This results in an abrupt change in trajectory, the animal being redirected towards the respective ‘selected’ target (Fig. 1C; see also Fig. S3A for the same phenomenon occurring for a wide range of starting positions).

**Fig. 1.**
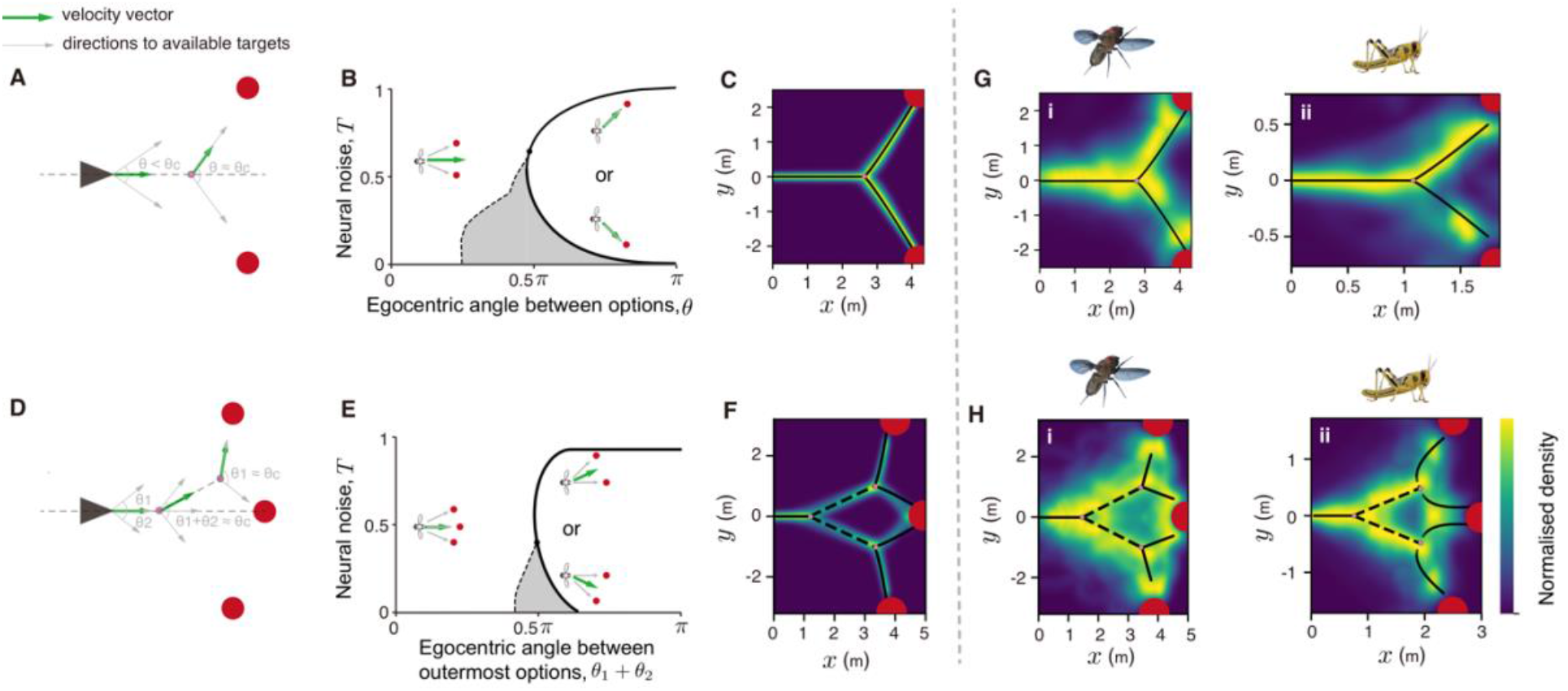
Geometrical principles of two-choice and three-choice decision-making. (A) Schematic of the binary decision-making experiments. This simplified representation shows that a sharp transition in the animal’s direction of travel is expected near a critical angle, *θ*_*c*_. (B) A phase diagram describing the ‘critical’ transition exhibited while moving from compromise to decision between two options in space. The shaded area (also in E) represents the region in parameter space where both the compromise, and the decision solutions exist. (C) Density plot showing trajectories predicted by the neural model in a two-choice context. The axes represent *x* − and *y* −coordinates in Euclidean space. The black line (also in G) presents a piecewise phase-transition function fit to the bifurcation. (D) Schematic of three-choice decision-making experiments, where the central target is on the angle bisector of the angle subtended by the other two targets. (E) A phase diagram describing the first ‘critical’ transition when the individual chooses among three options. Once the individual eliminates one of the outermost targets, it can decide between the two remaining options, similar to the two-choice phase diagram described in B. (F) Theoretical predictions for decision-making in a three-choice context. The dashed line (also in H) is the bisector of the angle subtended by center target and the corresponding side target on the first bifurcation point. See Table S1 for parameters used in C and F. (G) and (H) Density plots from experiments conducted with flies and locusts choosing among two and three options, respectively. Note that the density plots presented here are for the non-direct tracks, which constitute the majority type of trajectory adopted by both flies and locusts (Figs. S11 and S12). However, our conclusions do not differ if we use all, unfiltered, data (Figs. S11G,N and S12I,R).

Our model therefore predicts that despite the fact that the egocentric geometrical relationship between the animal and the targets changes continuously, upon approaching the targets, there exists a location whereby a further, very small, increase in angular difference between the targets will result in a sudden change in system (neural) dynamics, and consequently in motion, and thus decision-making. Such spatio-temporal dynamics do not occur if individuals were to simply integrate noisy vectorial information or choose their travel direction from a summed distribution of the location of targets in their sensory field (19), points we will return to later.

In numerical analysis of our model we find that irrespective of starting position, as the animal reaches the respective angle in space, it will relatively suddenly select one of the options (Fig. S3A). While the specific angular difference at which this phenomenon occurs is dependent on neural tuning, *ν* (Fig. S3C), and the starting configuration (due to an interplay between the two timescales involved—for movement and for neural consensus, see Fig. S3B), it is always present as long as the neural noise, *T* remains below a critical firing rate, *T*_*c*_ (although even for *T* < *T*_*c*_, these bifurcations may be difficult to see for small values of *ν* due to inherent noise in real biological systems; see Fig. S4 for simulations where vectorial representations of targets include directional error).

To gain a deeper insight into the mechanism underlying the observed spatiotemporal dynamics, we constructed a mean-field approximation of our model (see SI Appendix) since this has the advantage of allowing us to conduct formal analyses of patterns realized in the simulated trajectories.

## Geometric principles of decision-making

The mean-field analysis of our model shows that below a critical level of neural noise, animals will adopt the average among options as they approach the targets, until a critical phase transition upon which the system spontaneously switches to deciding among the options (Figs. 1B and S5A). Thus despite varying in its exact location (Fig. 1B), the sudden transition observed is an inevitable consequence of the system dynamics and will always occur.

Such sudden transitions correspond to ‘bifurcations’ in the mathematical study of dynamical systems. A bifurcation is said to occur when a smooth change in an external parameter, in this case perceived angular difference between the options, causes a sudden qualitative change in the system’s behavior, here corresponding to a literal bifurcation (or branching) in physical space.

When dynamical systems undergo such a phase, or quasi-phase, transition they exhibit a remarkable universal property: close to the transition, at the “critical-point” or “tipping-point”, the system spontaneously becomes extremely sensitive to very small perturbations (e.g. to small differences in preference between options (20, 21)). This is true of both physical (e.g. magnetic (22)) and biotic (e.g. cellular (23, 24)) systems undergoing a phase transition. Correspondingly, we find that below a critical level of neural noise, the mean-field model exhibits a sudden increase in susceptibility as the animal approaches the critical point, immediately prior to the decision being made (Fig. S5A). This will not occur in previously-considered models where an animal is assumed to choose its direction of travel based on the summed distribution of targets in its sensory field, also known as probability density function, or PDF, sum-based models (19). Thus, as animals approach targets we predict they will pass through a window of space (corresponding to the critical angle for the respective geometry they are experiencing) in which their brain spontaneously becomes capable of discriminating between very small differences between options (e.g. a very small difference in neuronal activity being in ‘favor’ of one option; see Fig. S3D and SI Appendix for details). This highly-valuable property (for decision-making) is not built into the model, but is rather an emergent property of the inherent collective dynamics.

In many real biological systems, including the ones we consider here, the (neural) system size is typically not large enough to consider true phase transitions (which only occur for very large systems, as per the mean-field approximation), but rather ‘phase-transition-like’, or ‘quasi-phase transition’, behavior. Even though real biological systems are not necessarily close to the infinite size limit of the mean-field approximation, we see very similar dynamics for both small and large system sizes (Fig. S6).

## Decision-making beyond two options

While the majority of decision-making studies consider only two options (due to both theoretical and experimental tractability (14, 25, 26)), animals moving in real space frequently encounter a greater number than this. Here we consider how animals will be expected to select among three, or more, options (possible targets) in space. First we begin with three identical options (*N*_1_ = *N*_2_ = *N*_3_) since this gives us the clearest insight into the relationship between motion and decision-making dynamics. Then we relax these assumptions and consider differences between options (Fig. S3E) as well as a greater number of options (Fig. 2). Note that we do not modify our model in any way prior to introducing these additional complexities.

**Fig. 2.**
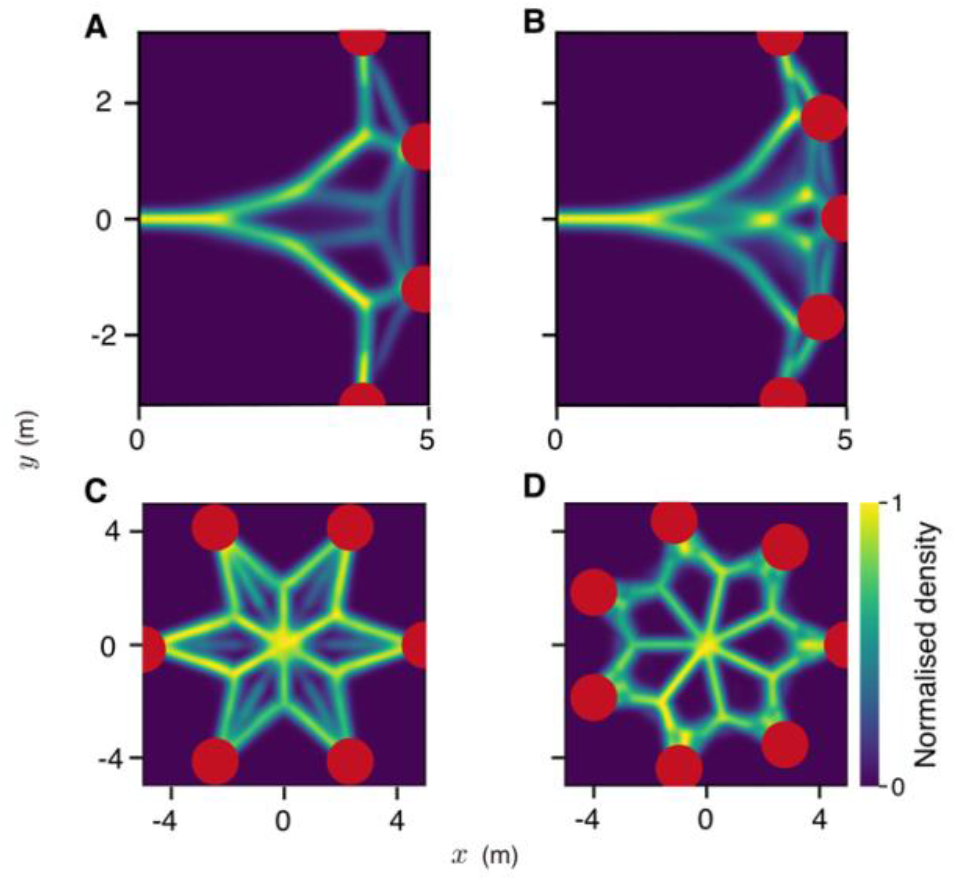
Decision-making for a larger number of targets. Density plots of simulated trajectories for four-(A), five-(B), six-(C) and seven-choice (D) decision-making when targets are placed equidistant and equiangular from the agent. Thee axes represent *x* − and *y* −coordinates in Euclidean space. Geometrical configurations are also varied to place the targets on the same side of the agent (A and B) or in radial symmetry (C and D). See Table S1 for parameters used in A–C. In D, all parameters used are identical except the system size *N* = 70.

Below *T*_*c*_ (see SI Appendix and Fig. S7 for considerations when *T* > *T*_*c*_), we once again find that the direction in which the animal moves is a function of the angular difference between the targets. When relatively far from the targets, it moves in the average of these three directions. Upon reaching a critical angular threshold between the leftmost and rightmost option (from the animal’s perspective), however, the neural system spontaneously eliminates one of them and the animal begins moving in the direction average between the two remaining options (Fig. 1D,E). It continues in this direction until a second critical angle is reached, and now the animal eliminates one of the two remaining options and moves towards the only remaining target (Figs. 1F and S5B). Thus we predict that the brain repeatedly breaks multi-choice decisions into a series of binary decisions in space-time. Such bifurcation dynamics are not captured in models of decision-making that do not include the required feedbacks, such as if individuals simply sum noisy vectors (or PDFs) to targets in their sensory field (19). For the case of three targets, vectors/votes to the leftmost option would tend to cancel those that favor the rightmost option, resulting in the selection of the central option, an issue we will return to later when considering collective animal behavior. Simulating a larger number of options (Fig. 2) and varying environmental geometries (Figs. S8 and S9) demonstrate the robustness of this mechanism in the face of environmental complexity and the more complex spatial dynamics that emerge as organisms undergo repeated bifurcations.

## Experimental tests of our predictions

Since the decision-process is predicted to be sequential, and dependent on the geometry with respect to the targets from an egocentric perspective, it should be possible to visualize it directly from the trajectories taken by animals when making spatial decisions. In this respect, our theoretical studies make a key testable prediction: if neural groups within the decision-making ensemble exhibit relatively local excitation, and long-range/global inhibition, we should observe bifurcations in the animals’ trajectories as they choose among identical options; and that if animals face three (or more) such options, then the complex decision task should be broken down to a series of binary decisions.

Since the geometrical principles revealed above are expected to be both robust and generic, we use immersive virtual reality (27) (Fig. S10) to test our predictions by investigating both two- and three-choice decision-making in three evolutionarily highly-divergent brains under ecologically-relevant scenarios: fruit flies (*Drosophila melanogaster*) and desert locusts (*Schistocerca gregaria*) deciding which among multiple vertical objects to approach (e.g. to perch), and zebrafish (*Danio rerio*) choosing with which conspecific(s) to school.

Like many other insects (28–31), fruit flies (32) and desert locusts (33) exhibit a natural tendency to orient and move towards high-contrast vertical features (potential landing sites or indicators of vegetation) in their environment. We exploit this tendency, presenting multiple identical black pillars as targets in an otherwise white environment. We record trajectories of our focal animals (solitary flies or locusts) as they choose to move towards one of these pillars, thus obtaining a behavioral readout of the decision-making process (see SI Appendix for experimental details; Figs. S11 and S12 show raw trajectories of flies and locusts respectively).

As predicted by our theory (Fig. 1B,C), we find that, in the two-choice case, most flies and locusts that choose one of the presented targets initially move in the average of the egocentric target directions until a critical angular difference (Fig. S13), at which point they select (randomly) one, or the other, option and move towards it (randomization test where *y* −coordinates between trajectories were swapped showed that the bifurcation fit to our experimental data was highly significant; *p <* 0.01 for both flies and locusts; Figs. 1G and S13). Here, we note that there may be multiple factors that affect the animals’ direction of movement. For example, it could be that animals repeatedly switch between fixating on each of the two options before reaching the critical angular difference, following which it selects one. However, quantification of their heading relative to the targets, and to the average direction between the targets (Fig. S13), finds no evidence for this; instead, prior to the bifurcation, both flies and locusts exhibit a heading towards the average of the egocentric target directions. In the three-choice case, the animals’ movements are also consistent with our theory; as predicted (Fig. 1E,F) they break the three-choice decision into two sequential binary decisions (*p* < 10^−4^ for both flies and locusts; Fig. 1H). For both animals, the observed angle of bifurcation (∼110° for flies and ∼90° for locusts) is much larger than their visual spatial resolution (∼8° and ∼2° for flies (34) and locusts (35, 36), respectively). We note ∼30% of animals in our experiments (both flies and locusts) did not exhibit the sequential bifurcations (see Figs. S11 and S12) described above, and instead moved directly towards one of the presented targets (Figs. S11 and S12). Such variability in response is expected in animals, and is consistent with recent work on the visual response of flies, which demonstrates a link between stochastic (non-heritable) variation in brain wiring within the visual system and strength of visual orientation response to a vertical stripe target (37). Furthermore, flies that experience high temperatures during development appear to exhibit a particularly strong orientation tendency, exhibiting the most direct paths to targets while flies that experience low developmental temperatures exhibit wandering paths to targets (38). In our model such differences can be accounted for by variation in directional tuning of the neural groups, with high directional tuning (low *ν*) being associated with a strong orientational response, and such individuals exhibiting direct tracks to targets from the outset (see Fig. S14).

A further, non-mutually exclusive, possibility, is that a subset of insects exhibit “handedness”. For example, in (39), it was shown that approximately 25% of *Drosophila* were either strongly left-biased or right-biased when moving on a Y-maze, and that these consistent differences among flies were similarly non-heritable. This experimental design did not assess whether a further subset are biased to go directly forwards if offered three directional choices (such as could occur in a hypothetical Ψ maze). In such cases, it is certainly possible that these intrinsic directional biases break symmetry (Fig. S3D,E), resulting in directed paths to different targets.

We note that individuals predisposed to exhibit direct paths to targets would be expected to make faster, yet less accurate, decisions, a prediction we plan to test in future studies.

Our zebrafish experiments consider spatial decision-making in a social context. We present virtual conspecifics (see SI Appendix for methodological details) that move back-and-forth in the arena parallel to each other as targets (Figs. 3A and S15A) and behave (Fig. S16), and are responded to (Fig. S17), in the same way as real fish. Because they are social, the real fish respond to these virtual fish by tending to follow at a (relatively) fixed distance behind them (Fig. S15E). Our data are best represented within this moving frame of reference (the virtual fish; Fig. S15). Theoretically we predict that for two virtual fish we should see a single bifurcation, where the real fish will suddenly switch from averaging the target directions to deciding among them (i.e. swimming predominantly with one of the virtual fish), as a function of increasing the lateral distance, *L*, between the virtual fish (Figs. 3B and S18; see SI Appendix for details of model implementation). The existence of this bifurcation is clearly seen in our experiments (Fig. 3C). When considering three moving virtual conspecifics, the model predicts that real fish will spontaneously break the three-choice decision to two binary decisions, and a comparison of the theoretical prediction and experimental results demonstrates this to be the case (c.f. Fig. 3E,F).

**Fig. 3.**
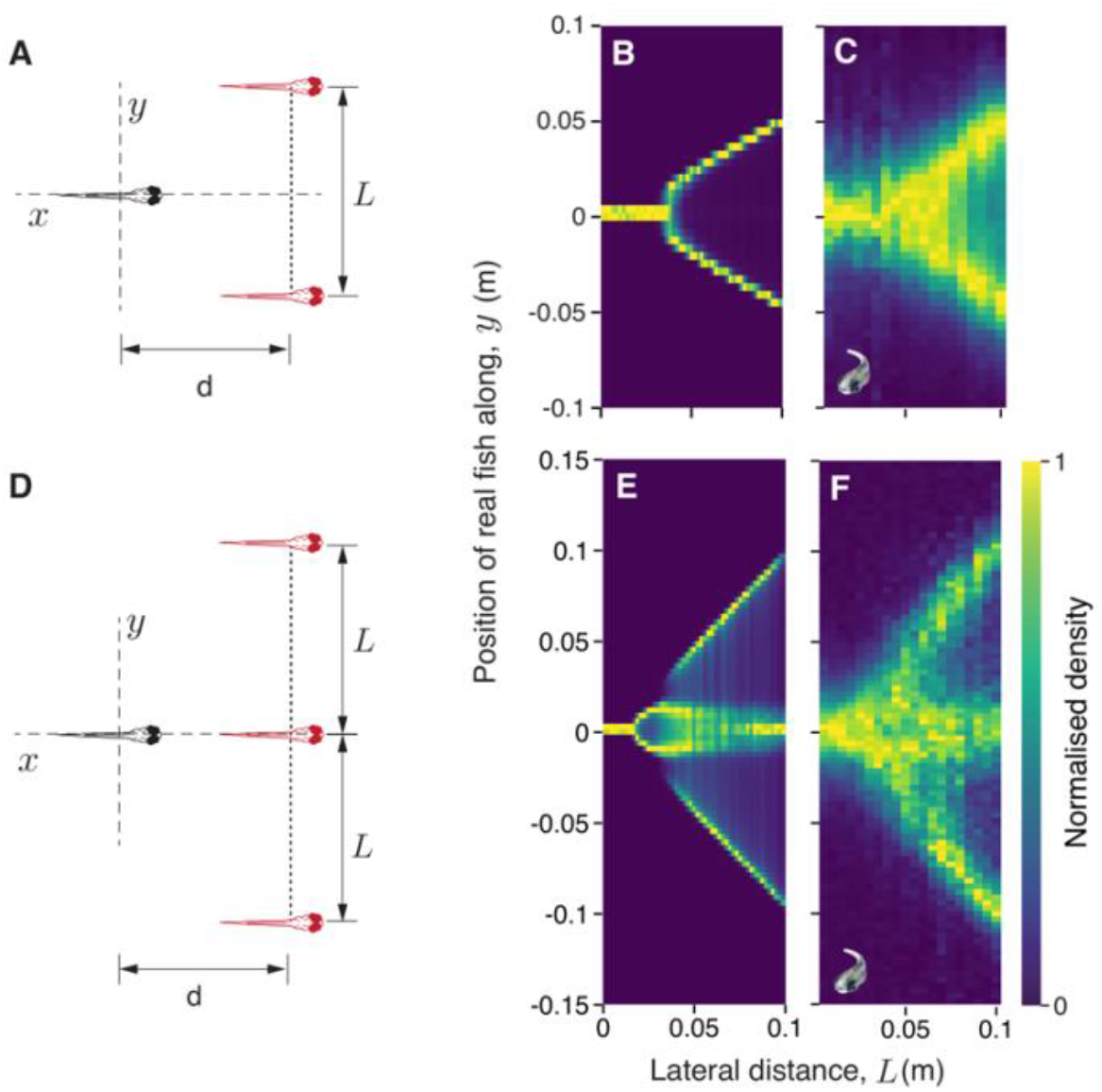
Decision-making in a moving frame-of-reference. (A) Schematic of the two-choice decision-making experiments conducted with larval zebrafish. In these experiments (also in the three-choice experiments depicted in D), the virtual fish swim parallel to each other while maintaining a fixed lateral distance, *L* between them. We only consider data where the real fish swims behind the virtual fish, i.e., it follows the virtual fish (see SI Appendix and Fig. S15 for details). (B) Normalized probability distribution (proportion of maximum) of simulated positions of an agent following two moving targets, and corresponding experiments (C) conducted with larval zebrafish following two virtual conspecifics. (D) Schematic representation of the three-choice decision-making experiments. (E) Normalized probability distributions of simulated positions of an agent following three moving targets, and corresponding experiments (F) conducted with larval zebrafish following three virtual conspecifics. See Table S1 for model parameters used in B and E.

We also test predictions under conditions where there is an asymmetric geometry whereby two fish swim closer to each other than the central one does to the third fish (Fig. 4A). As predicted by our theory (Fig. 4B), the real fish tends to swim between the two closely-associated fish, or close to the third, more distant, fish (Fig. 4B). Note that, also as predicted, the real fish spends a similar amount of time in each of the two locations.

**Fig. 4.**
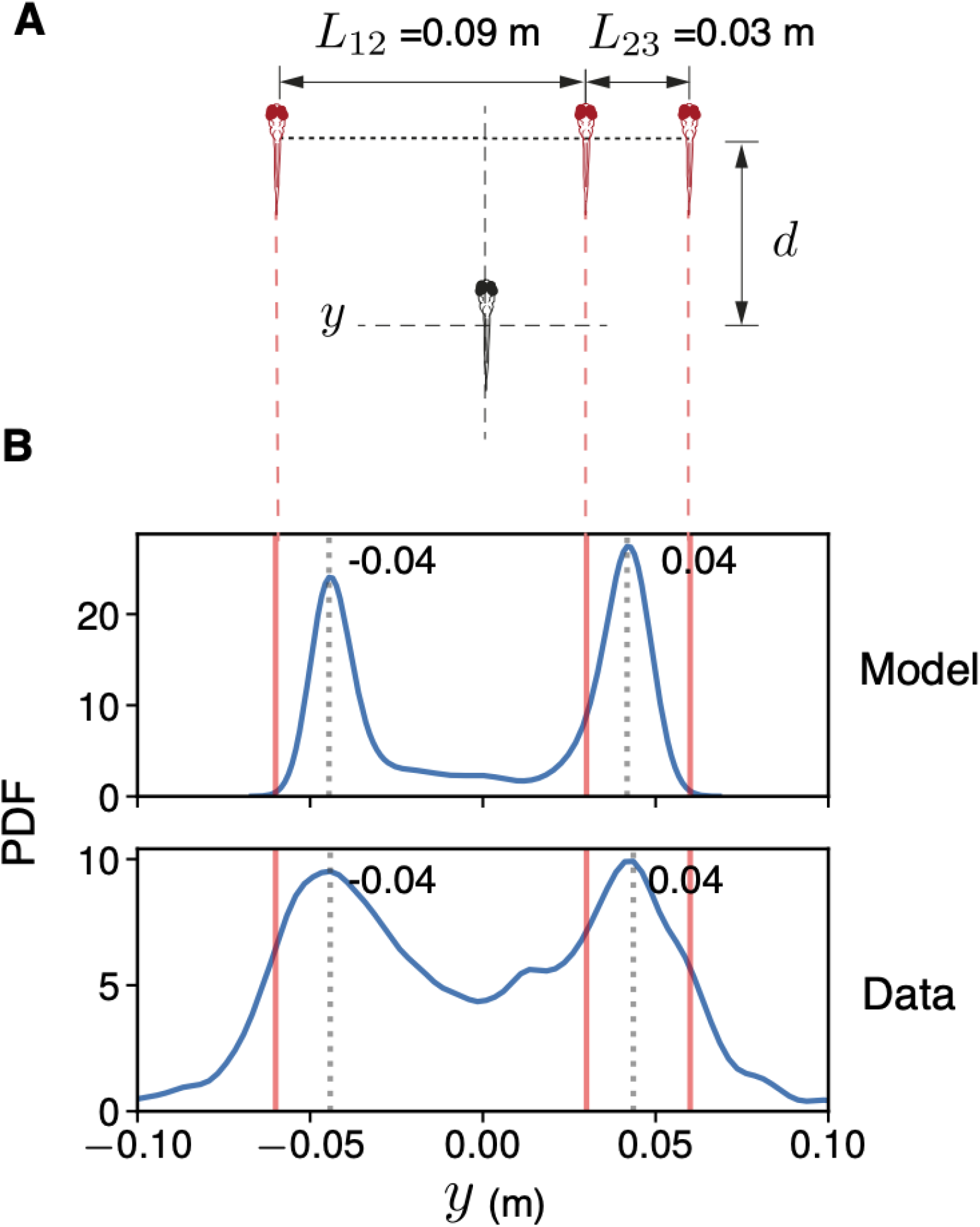
Decision-making with the targets in an asymmetric geometry. (A) Schematic of the asymmetric choice test presented to larval zebrafish. In these experiments, the virtual fish swim parallel to each other while maintaining a fixed lateral distance, *L* between them. To create asymmetry in the geometry, the center fish swims closer to one of the side fish than the other (*L*_12_ = 0.09 m and *L*_23_ = 0.03 m). (B) The upper panel shows the probability density function of simulated positions of an agent following three moving targets in an asymmetric geometry corresponding to the experiments. The simulated agent occupies a position of *y* = ±0.04 m while following the targets (ν = 0.7; *σ*_*θ*_ = 0.3). The lower panel shows the probability density function of the position of the real fish along the axis perpendicular to its direction of motion. As predicted by our model, the real fish considers the two virtual conspecifics closer to each other as a single target and adopts one of two positions behind the virtual fish.

Although detailed models considering the specifics of each system would be expected to provide additional quantitative fits (at the expense of losing some degree of generality and analytical tractability), our results are broadly independent of the model implementation details. Thus, we find that the key predictions of our model are validated in fruit flies, desert locusts and larval zebrafish in distinct, yet ecologically relevant contexts.

## Model features that determine network behavior

There are key features that are essential to produce the bifurcation patterns observed in our data i.e. for any decision-making system to break multi-choice decisions to a series of binary decisions.

- Feedback processes that provide the system directional persistence, and drive such bifurcations, are crucial to exhibit the observed spatio-temporal dynamics. In the neural system, this is present in the form of local excitation and long-range/global inhibition (7, 16, 17). However, as shown in our model of collective animal behavior below, we expect that similar dynamics will be observed if the necessary feedbacks are also incorporated into other models of decision-making, such as to PDF-sum-based models, for example (19).
- Observing similar decision dynamics requires a recursive (embodied) interplay between neural dynamics, and motion in continuous space. Here, the animal’s geometrical relationship with the targets changes as it moves through physical space. Since neural interactions depend on this changing relationship, space provides a continuous variable by which the individual traverses the time-varying landscape of neural firing rates.

These essential features, along with the observed animal trajectories in the two-choice context, are reminiscent of collective decision-making in animal groups (models (40–44), fish schools (45), bird flocks (46) and baboon troops (25)). Below, we consider an established model of collective decision-making (40) to draw links between these two scales of biological organization—decision-making in the brain, and decision-making in animal groups.

## A link to collective decision-making

In order to draw a link between individual decision-making and collective decision-making in animal groups, we consider an animal group with equal number of individuals exhibiting preference for each target (see SI Appendix for methodological details). A long-standing approach in the study of animal collectives is to consider them integrating vectorial information from neighbors (47, 48), and there are a great number of publications of such “flocking”, “schooling” or “herding” behaviors (47–49). Individuals within groups may also have preferences to reconcile this local vector-averaging with goal-oriented behavior, such as a desired direction of travel (40, 45), and such models have made effective predictions regarding how the number of individuals with a common desired direction of travel influences the accuracy of group motion towards targets (25) and how the weighting of the internal ‘goal-oriented’ vector representing the desired direction of travel, influences the capacity and accuracy for individuals to act as leaders and to influence the directon taken by the group as a whole (50).

We demonstrate here, however, that while ubiquitous, such models of collective animal behavior fail to account for the known capability for animal groups to make decisions among spatially discrete targets (see Fig. S19A,B). To do so, it is essential that the necessary feedbacks, as described above for collective decision-making among neurons, are incorporated. While these feedbacks are inherent to our neural model, they can also be included in other models in the form of social interactions, or in the animals’ response to their environment (51).

For example, one way feedback can be introduced here is via ‘informed’ individuals (those with a desired direction of travel) associating with ‘uninformed’ or ‘unbiased’ individuals (individuals that exhibit social interactions but have no specific desired direction of travel) (40, 45); ‘uninformed’ individuals are effectively recruitable by those with a desired direction of travel (providing local positive feedback), but are also in finite supply, creating what is effectively a competition among informed subsets that differ in their preferred direction of travel (a form of longer-range inhibition between informed subsets). However, because ‘uninformed’ individuals tend to average the direction of all ‘informed’ individuals that recruit them, we find that this type of feedback functions more as a social glue, and is only able to explain bifurcations when the group is choosing between two options. In a decision-making context with three options, this type of feedback, alone, results in the group almost always moving towards the central target (Fig. S19D).

A means of resolving this issue is for individuals to change the strength of their goal-orientedness as a function of their experienced travel direction; for example, individuals that find themselves consistently moving in a (group) direction that differs from their preferred target direction could weaken the strength of their preference over time (a form of forgetting/negative feedback, effectively resulting in long-range/global inhibition; and once this preference is lost, they will tend to spontaneously reinforce the majority-selected direction (45), a form of positive feedback). We find that this biologically-plausible mechanism (40) will allow individuals within the group to recover the capability to come to consensus even in the absence of uninformed individuals (Fig. 5), and for a greater number of options than two (Fig. 5B).

**Fig. 5.**
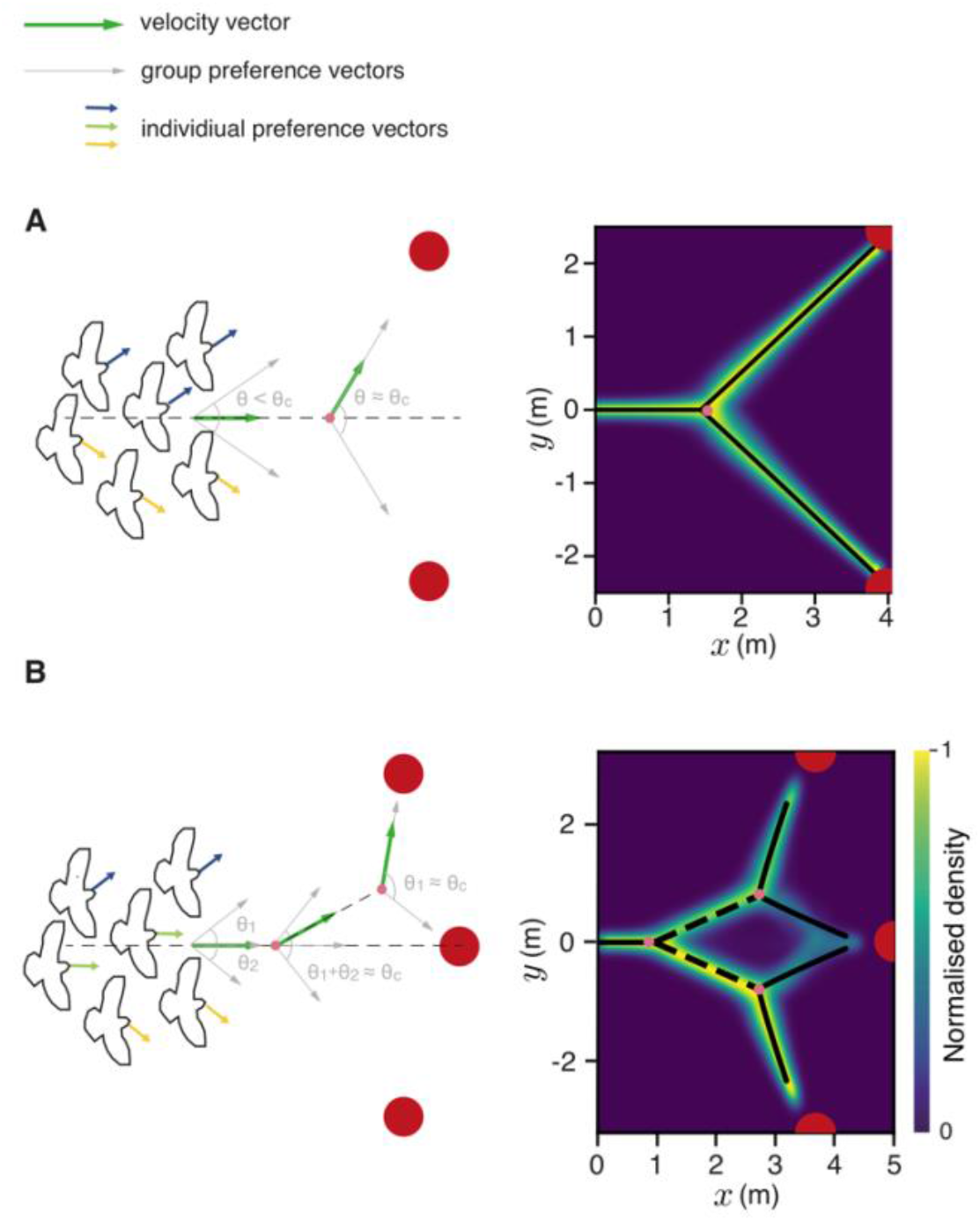
Consensus decision-making in simulations of animal groups follow the same geometrical principles. Results for two-(A) and three-choice (B) decision-making in a model of animal collectives. The density plots show trajectories adopted by the centroid of the animal group for 500 replicate simulations where the groups don’t split. The axes represent *x* – and *y*−coordinates in Euclidean space. The black lines show a piecewise phase-transition function fit to the trajectories. For the three-choice case (B), the dashed line is the bisector of the angle subtended by center target and the corresponding side target on the first bifurcation point. See Table S2 for parameters used.

Despite considerable differences in details between this model and that of neural dynamics described above, with the former involving individual components that change neighbor-relationships over time and where inhibition emerges from a different biological process, the predictions regarding motion during decision-making are extremely similar (c.f. Fig. 5 and Fig. 1 for a comparison between predictions for animal groups and neural groups, respectively). Thus, we find that similar principles may underlie spatial decision-making across multiple scales of biological organization. Furthermore by presenting social interactions in a decision-making context, our zebrafish experiments elucidate the neural basis of schooling allowing us to glean insights across three scales of biological organization—from neural dynamics to individual decisions, and from individual decisions to collective movement.

## Conclusions

We demonstrate that, across taxa and contexts, explicitly considering the time-varying geometry during spatial decision-making provides new insights that are essential to understand how, and why, animals move the way they do. The features revealed here are highly robust, and we predict that they occur in decision-making processes across various scales of biological organization, from individuals to animal collectives (see Figs. 5 and S19, and SI Appendix), suggesting they are fundamental features of spatiotemporal computation.

## Materials and Methods

We construct a simple, spatially-explicit model of neural decision-making to study how the brain reduces choice in the presence of numerous spatial options (adapted from (52)). Theoretical predictions obtained were then tested experimentally by exposing invertebrate (fruit flies and desert locusts) and vertebrate systems (zebrafish) to spatial choice tests in virtual reality. To identify unifying principles of spatiotemporal computation across scales of biological organisation, we also reproduce the obtained decision-making patterns with an established model of collective decision-making in animal groups.

### Neural decision-making model

We construct a computational model of neural decision-making that takes in a representation of directions to the different targets as input, and outputs a collective vectorial representation of the agent’s future velocity (adapted from (52)). This provides us explicit predictions for animal trajectories, allows us to determine which target is reached in each realization of the simulation, and facilitates direct comparison with experimental tests. Our model is within the class of widely-employed neural ring-attractor models (see SI Appendix), which like neural field models (53, 54), and attractor network models more generally (15, 16, 55), consider the collective firing activity of the neurons, or the firing rate, as opposed to the microscopic state of each firing neuron.

In our model, the brain is composed of individual components, called “spins”, that, collectively, as a “spin system”, represent neural activity. Spin systems, which have been long-studied in physics due to their ability to give insight into a wide range of collective phenomena, from magnetic to quantum systems (56), were first introduced in the study of neurobiology by Hopfield in a landmark paper (57) that provided considerable insights into principles underlying unsupervised learning and associative memory. In its simplest (and most common) formulation, as in Hopfield networks, a spin system is comprised of entities, spins, that can each be in state 0 or 1, or in the terminology of physics either ‘up’ or ‘down’. Spin systems have consistently provided deep insights into complex collective phenomena, from spin and molecular systems, to neural systems, undergoing phase transitions (58, 59) (see SI Appendix for details and discussion).

Here, the animal’s brain is characterized by a system of $N$ spins. Each spin $i$ encodes direction to one of the presented goals $\hat{p}^_i$, and exists in one of two states: $\sigma_i=0$ or $\sigma_i=1$. We do not imply that a spin is equivalent to a neuron, but rather, as we show via a mathematical derivation, that the collective properties of interacting spins in our model is equivalent to the firing rate in the neural ring attractor model (see SI Appendix for details). Consequently, we refer to the individual components with which we model our system as “spins”, and “neural activity” as a term to represent this “firing rate” equivalent. The energy of the system (for any given configuration) is given by its Hamiltonian, *H*.

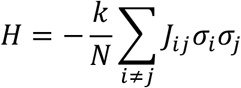

where, *k* is the number of options available to the individual and *J*_*ij*_ is the interaction strength between neurons *i* and *j*. Here, *J*_*ij*_ is given by

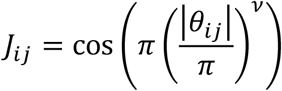

where, *θ*_*ij*_ is the angle between preferred directions of neurons *i* and *j*, and *ν* represents the neural tuning parameter. For *ν* = 1, the interactions become “cosine-shaped” *J*_*ij*_ = cos(*θ*_*ij*_), and the network has a Euclidean representation of space (Fig. S1). For *ν* < 1, the network has more local excitation and encodes space in a non-Euclidean manner (Fig. S1). System dynamics are implemented by energy minimization using the Metropolis-Hastings algorithm (similar to other Ising spin models) and the agent then moves with a velocity 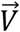 determined by the normalized sum of goal vectors 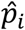 of all active neurons.

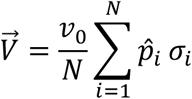

where *v*_0_ is the proportionality constant. The goal vector 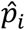 now points from the agent’s updated location to the neuron’s preferred goal with directional noise chosen from a circularly wrapped Gaussian distribution centered at 0 with a standard deviation *σ*_*e*_. As in the mean-field approximation of the model, the timescale of movement (defined by the typical time to reach the target) in the numerical simulations was set to be much greater than the timescale of neural firing (the typical time between two consecutive changes in the neural states *σ*_*i*_).

### Collective decision-making model

We reproduce results from our neural decision-making model in a model that describes spatial decision-making at a different scale of biological organization (refer (40) for methodological details). To highlight the features that are key to producing the observed bifurcation patterns, we run simulations with and without feedback on the strength of goal-orientedness of individuals.

### Fly virtual reality experiments

All experiments were conducted on 3- to 5-day old female wild-type CS strain *Drosophila melanogaster* raised at 26°C on a 12 hr light, 12 hr dark cycle. Experiments were conducted in a flyVR setup procured from loopbio GmbH. 60 tethered *Drosophila melanogaster* were exposed to either a two-choice or a three-choice decision task (30 and 30 individuals, respectively) in the virtual reality environment. Each experimental trial lasted 15 min where flies were exposed to five sets of stimuli—three experimental sets and two control sets. The experimental stimuli sets consisted of two or three black cylinders (depending on the experimental condition) that were presented to the animal in an otherwise white environment. A control stimulus with a single pillar was presented before and after the experimental conditions. We rotated all trajectories such that the *x* −axis points from the origin, to the centre of mass of the targets. To visualise trajectories in the various experimental conditions, we created time-normalised (proportion of maximum across a sliding time window) density maps. We then folded the data about the line of symmetry, *y* = 0 and applied a density threshold to the time-normalised density map. A piecewise phase transition function was then fit to quantify the bifurcation.

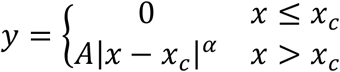

where *x*_*c*_ is the critical point, *α* is the critical exponent, and *A* is the proportionality constant. We also performed randomisation tests for each bifurcation where we conducted the exact fit procedure described above to data where the trajectories were randomised by keeping the *x*-coordinates, and swapping the *y*-coordinates with values from other random events. Randomizations show that the resultant fit to our experimental data were highly significant (*p <* 0.01 for binary choice and *p <* 10^−4^ for the three-choice case).

Based on the amount of time it took flies to reach one of the available targets, we also classified individual fly tracks into one of two categories—direct tracks and non-direct tracks (60) (see Fig. S11A,H for details). In our model, the direct tracks were also accounted for by varying the directional tuning of spins. A high neural tuning (low *ν*) results in more directed tracks (Fig. S14).

### Locust virtual reality experiments

All experiments were conducted on 156 instar 5 desert locusts (*Schistocerca gregaria*; 57 individuals for two-choice and 99 individuals for three-choice experiments, respectively) raised in the Animal Research Facility of the University of Konstanz. Based on our filtering criteria, 122 out of the 156 locusts were used in our analyses. Experiments were conducted in a locustVR setup procured from loopbio GmbH (27). The experimental procedure was identical to the one described above for flies, except now, each experimental trial lasted 48 min—three experimental sets (12 min each) and two control sets (6 min each). Analyzing bifurcations in locust trajectories using the same methods described above showed that the resultant bifurcations fit to our experimental data were highly significant (*p <* 0.01 for binary choice and *p <* 10^−4^ for the three-choice case).

Similar to the flies, the locust trajectories were also classified as direct, or non-direct tracks. However, because the locustVR system allowed the animals to stop and reconsider movement during the decision-making process, we added an additional category to classification of individual locust tracks *viz*. the wandering tracks (see Fig. S12A,J for details).

### Fish virtual reality experiments

All experiments were conducted on 1 cm ± 0.1 cm long zebrafish (*Danio rerio*) of age 24 to 26 days post-fertilisation raised in a room at 28 °C on a 16 hr light, 8 hr dark cycle. 440 fish were tested in total. Of these, 198 fish were exposed to decision-making with two virtual targets, 39 fish were exposed to decision-making with three equidistant virtual targets, and 50 fish were exposed to decision-making with three targets in asymmetric geometry (see SI Appendix for more details). Experiments were conducted in a fishVR setup procured from loopbio GmbH (refer (27) for details). Once a fish was introduced in the arena, it was given 20 min to acclimatize to the environment. This was followed by a 10 min control where it was presented a single virtual conspecific circling the arena in a circle of radius 8 cm. After this, for experiments in symmetric geometries, the real fish was exposed to choice experiments that lasted 90 min with the virtual fish initialized with random lateral distances between them and random swim direction. To visualize the bifurcations, we normalized (proportion of maximum) and stacked the marginal distributions along the direction of the virtual fish’s motion for various lateral distances. For experiments in asymmetric geometries, the real fish was exposed to choice experiments where distance between the center virtual fish and its closer neighbor was 0.03 m and its distance to the other neighbor was 0.09 m (Fig. 4). All experiments were conducted in accordance with the animal ethics permit approved by Regierungspräsidium Freiburg, G-17/170.

## Acknowledgements

We thank all members of the Department of Collective Behaviour who assisted with the project: Renaud Bastien for modelling discussions and help setting up the VR experiments, Guy Amichay for providing control data of two real fish swimming together, and Paul Szyszka for showing V.H.S. how to tether flies. We thank the ‘Itai Cohen Lab’ for the fruit fly image used in Fig. \ref{fig:insects}, Jitin Thomas for help with visualizations in Unity and Andreas Poehlmann, John Stowers and Max Hofbauer from loopbio GmbH for technical support with the VR systems. V.H.S., L.L., B.R.S. and I.D.C. are also grateful to the animal care at the University of Konstanz including Christine Bauer, Jayme Weglarski and Dominique Leo for help in conducting the experiments. They also acknowledge the efforts of the scientific and technical staff at the University of Konstanz including Michael Mende, Markus Miller, Mäggi Hieber Ruiz and Daniel Piechowski. V.H.S. acknowledges the International Max Planck Research School (IMPRS) for Organismal Biology for the graduate school community and access to courses and resources. I.D.C. acknowledges support from the NSF (IOS-1355061), the Office of Naval Research grant (ONR, N00014-19-1-2556), the Struktur-und Innovationsfonds für die Forschung of the State of Baden-Württemberg, the Deutsche Forschungsgemeinschaft (DFG, German Research Foundation) under Germany’s Excellence Strategy-EXC 2117-422037984 and the Max Planck Society. N.S.G. is the incumbent of the Lee and William Abramowitz Professorial Chair of Biophysics and acknowledges support by the Minerva Foundation (grant no. 712601). M.N acknowledges support from the Eötvös Loránd Research Network (a grant to the MTA-ELTE ‘Lendület’ Collective Behaviour Research Group, grant number 95152, and MTA-ELTE Statistical and Biological Research Group) and Eötvös Loránd University. Additionally, all authors thank two anonymous referees for their constructive comments during the review process.

## Author contributions

V.H.S. and I.D.C. designed the study; V.H.S., D.G., T.S., N.S.G. and I.D.C. constructed the model; D.G. and N.S.G. constructed the mean-field approximation; V.H.S. and I.D.C. designed the fly experiments; V.H.S. conducted these experiments and analyzed the data with L.L. and M.N.; V.H.S., B.R.S. and I.D.C. designed the locust experiments; B.R.S. conducted these experiments and V.H.S. analyzed the data with M.N.; L.L. and I.D.C. designed the fish experiments; L.L. conducted these experiments and analyzed this data with V.H.S. and M.N.; V.H.S. and I.D.C. drafted the manuscript with significant contributions from all authors.

## Competing interests

The authors declare that they have no competing interests.

## Notes

### Competing Interest Statement

The authors have declared no competing interest.

### Summary of Updates

Updated manuscript (based on reviewer comments)

