## Supplementary Information for "The geometry of decision-making"

### Materials and Methods

#### 1. Neural decision-making model

**1.1 Overview.** We construct a computational model of neural decision-making that takes in a representation of directions to the different targets as input, and outputs a collective vectorial representation of the agent's future velocity (adapted from (1)). This provides us explicit predictions for animal trajectories, allows us to determine which target is reached in each realization of the simulation, and facilitates direct comparison with experimental tests. Our model is within the class of widely-employed neural ring-attractor models (see section 1.8, below), which like neural field models (2, 3), and attractor network models more generally (4–6), consider the collective firing activity of the neurons, or the firing rate, as opposed to the microscopic state of each firing neuron.

Neural activity/firing rate models are very widely employed in neuroscience, having been applied to almost every area of study, from perception (7–9) to memory (10) and decision-making (11, 12). As has been found in the study of dynamical systems and statistical physics more generally, the governing equations in such systems often do not depend on microscopic details (13, 14). For example, just as it is not necessary to model molecular dynamics to simulate an individual neuron, many possible details of the individual components of a network do not influence the general collective dynamics exhibited, thus allowing for abstraction and a focus on revealing and understanding general principles (for an example of work that shows there exists an exact correspondence between physiological properties such as the mean firing rate and membrane potential within the network and low-dimensional dynamical properties, see (15)).

In direct relation to the present study, it has been established that there are ring-attractor dynamics in the *Drosophila* brain (16), and further, that the dynamics of this neural system appear to be highly robust to microscopic details. For example, (6) developed a detailed, physiologically-parameterized model, incorporating spiking neurons, with connectivity derived from microscopy of detailed neuronal morphologies in fruit flies. They found that their model behaved like a ring-attractor under a wide range of parameters, and that the ring-attractor properties were highly robust to physiological details, such as variation in synaptic weights. They concluded that "the ring-attractor computation is a robust output of this circuit, apparently arising from its high-level network properties (topological configuration, local excitation and long-range inhibition) rather than fine-scale biological detail". Further studies have shown that, in general, the attractor states of ring-attractor neural dynamics can be described by a low-level family of functions and thus that the solutions do not require, or depend on, explicit simulation of the neuronal dynamics (see (17) for details).

This is not to say that models accounting for individual neuronal properties and connectivity will not also be of importance. As we emphasise in our main manuscript, it is likely that detailed models will likely be able to better quantitatively fit specific systems. As (6) suggest, such models may prove helpful in understanding how thermogenetic and optogenetic manipulations to that specific network could impact behavior of specific organisms; but this will come at the cost of analytic tractability, and thus a weakened ability identify key principles that transcend system specifics. Thus, both (non-mutually-exclusive) approaches to modelling neural systems are valuable, depending on the purpose of scientific study.

**1.2 Framework.** In our model, the brain is composed of individual components, called "spins", that, collectively, as a "spin system", represent neural activity. Spin systems, which have been long-studied in physics due to their ability to give insight into a wide range of collective phenomena, from magnetic (18) to quantum (19) systems, were first introduced in the study of neurobiology by Hopfield in a landmark paper (20) that provided considerable insights into principles underlying unsupervised learning and associative memory. In its simplest (and most common) formulation, as in Hopfield networks, a spin system is comprised of entities, spins, that can each be in state 0 or 1, or in the terminology of physics either 'up' or 'down'. While at first appearing an almost absurd oversimplification of reality, spin systems have proven themselves by providing some of the deepest understanding of complex collective phenomena, and especially the universal properties, regardless be they spins, iron atoms or water molecules, undergoing phase transitions (21). More recently, it has since been shown, via analysis of real spiking neural networks, that such models are not only consistent with experimental data across scales, but that this similarity is not just an analogy or metaphor (see (22) for details of their system and analysis).

In our case, we do not imply that a spin is equivalent to a neuron, but rather, as we show via a mathematical derivation below, that the collective properties of interacting spins in our model is equivalent to the firing rate in the neural ring attractor model (see section 1.8 For details). Consequently, we refer to the individual components with which we model our system as "spins", and "neural activity" as a term to represent this "firing rate" equivalent. Our model is deliberately simple, our intention being to make robust predictions that transcend system specifics, and to establish the causal nature of the properties seen via mathematical analysis. Thus, a benefit of our chosen formulation is that we can take advantage of its analytic tractability, as well as the large body of knowledge regarding collective properties of spin systems, including a well-defined notion of stochasticity (noise), while also ensuring it is within the class of ring-attractor models that are so widely-employed in neuroscience.

Here, the brain is characterized by a recurrent neural network composed of $N$ spins. Each spin $i$ encodes direction to one of the presented goals $\hat{p}_i$, and exists in one of two states: $\sigma_i = 0$ or $\sigma_i = 1$. Since each spin exists in one of two states, there are $2^N$ possible system configurations. Energy of the system (for any given configuration) is given by its Hamiltonian, $H$.

$$H = -\frac{k}{N}\sum_{i\neq j} J_{ij}\sigma_i\sigma_j \quad [1]$$

where, $k$ is the number of options available to the animal and $J_{ij}$ is the interaction strength between spins $i$ and $j$. A positive $J_{ij}$ indicates an excitatory interaction between spin $i$ and spin $j$ while a negative $J_{ij}$ indicates an inhibitory interaction. Here, we assume that interactions are excitatory when spins encode a similar directional preference, and inhibitory when they encode conflicting directional preferences. This captures both explicit ring-attractor networks, with local excitation and long-range/global inhibition (as found in fruit flies, and other insects (16)), and computation among distributed competing neural groups (as in the mammalian brain (23)). The locality of excitatory interactions encoded by $J_{ij}$, or directional tuning of the spins is given by the tuning parameter $\nu$. Here, $J_{ij}$ is given by

$$J_{ij} = \cos\left(\pi\left(\frac{|\theta_{ij}|}{\pi}\right)^{\nu}\right) \quad [2]$$

where, $\theta_{ij}$ is the angle between preferred directions of spins $i$ and $j$, and $\nu$ represents the neural tuning parameter. For $\nu = 1$, the interactions become "cosine-shaped" $J_{ij} = \cos(\theta_{ij})$, and the network has a Euclidean representation of space (Fig. S1). For $\nu < 1$, the network has more local excitation and encodes space in a non-Euclidean manner (Fig. S1). For sake of simplicity, we assume a fully-connected network. At each timestep, energy of the system $H$ is minimized using the Metropolis-Hastings algorithm i.e. a change in neural state $\sigma_i$ is dependent on the change of energy ($\Delta H$) that accompanies it.

$$P^{(i)}_{1\to 0/0\to 1} = \begin{cases} \exp(-\Delta H/T) & \Delta H \geq 0 \\ 1 & \Delta H < 0 \end{cases} \quad [3]$$

where $P^{(i)}_{1\to 0/0\to 1}$ is the probability that a spin switches its state and $\Delta H = H_2 - H_1$ where $H_1$ is the energy of the system before the spin changes its state and $H_2$ is its energy after the change in state. This is akin to other Ising spin models where the temperature parameter $T$ is interpreted here as neural noise. The agent then moves with a velocity $\vec{V}$ determined by the normalized sum of goal vectors $\hat{p}_i$ of all active spins.

$$\vec{V} = \frac{v_0}{N}\sum_{i=1}^{N} \hat{p}_i\sigma_i \quad [4]$$

where $v_0$ is the proportionality constant. The agent moves along $\vec{V}$ and spins update their goal vector $\hat{p}_i$ to reflect the agent's movement. The goal vector $\hat{p}_i$ now points from the agent's updated location to the spin's preferred goal with directional noise chosen from a circularly wrapped Gaussian distribution centered at 0 with a standard deviation $\sigma_e$.

**1.3 Simulations.** We simulated an agent with sixty spins that is tasked with decision-making in an environment containing multiple (two–seven) targets (using seventy spins for the seven target case). Each spin was assigned a preference for one of the available targets which determined its goal vector $\hat{p}_i$. For each simulation run, the agent was reinitialized at (0, 0) and the targets were set at a distance of 5 units, corresponding to 5 m in the fly experiments. In the two-target case, the targets were separated by 60° putting them at (4.33, -2.5) and (4.33, 2.5) respectively. In the three-target case, successive targets were initialized to be 40° apart putting them at (3.83, -3.21), (5, 0) and (3.83, 3.21) respectively. We ran 500 replicate simulations for each condition.

To demonstrate sensitivity of this algorithm, we examine the effect of varying the proportion of spins pointing to a target, on the probability that it is chosen (Fig. S3D and E). The slope of the sigmoid here indicates how a small change in the proportion of spins encoding a target is amplified to maximize decision accuracy. We also vary the agent's starting position, distance to the target, and the neural tuning $\nu$ to illustrate the effect these parameters on the bifurcation angle (Fig. S3A-C).

Finally, we also simulate a scenario where the agent is restricted in motion only allowing movement in the direction perpendicular to direction to the targets. These simulations are meant to mimic a social decision-making scenario where the animal chooses to follow one of several moving conspecifics at a fixed distance behind them (see Fig. 3 in main text). Once again, the agent was initialized at (0, 0) while the targets were placed at a fixed distance d along the $x-$axis. We varied lateral distance between the targets such that their centroid was always at ($d$, 0) and recorded the agent's trajectory along the $y-$axis. Corresponding experiments were conducted with larval zebrafish (see Section 5 for experimental details).

**1.4 Parameter choice.** Our model is composed of two 'free' parameters, the neural noise parameter ($T$) and the neural tuning parameter ($\nu$). Here, we will discuss the values chosen for these parameters and their biological interpretation.

***1.4.1 Neural noise.*** The neural noise parameter $T$ represents the intrinsic spontaneous activity, a form of noise, present in neural systems, or spontaneous flipping of spins in our model. From the phase diagrams (see Fig. 1 in main text), it is evident that the animal will spontaneously transition from averaging direction to all available options, to eliminating one among the remaining options, so long as the brain is poised below a critical level of noise, $T < T_c$ (Fig. 1, B and E in main text and Fig. S7, A and C). Here, we will address behavior of the system when $T > T_c$ for decision-making in both the two- and the three-choice case. In the two-choice case, the agent consistently moves towards the average of the two target directions. However, any movement in the direction perpendicular to this can be described by a diffusive process (Fig. S7B). For the three-choice case, we observe that when $T$ exceeds $T_c$, geometrical relationship between options creates a strong bias towards central targets (Fig. S7D). This prevents the animal from exploring the different targets (of equal value) with a similar frequency—as is indicated by our experimental data where ratio of visiting a side target to visiting the center target is 0.82 for flies and 1.23 for locusts (this ratio would be 1 if all targets were visited equally). Thus, in order to maximize decision-accuracy in the presence of an arbitrary number of targets, biological systems (brains) must be poised below a critical level of neural noise, i.e. $T$ must be less than $T_c$.

***1.4.2 Neural tuning.*** The bifurcation angle $\theta_c$ increases monotonically with the neural tuning parameter, $\nu$ (Fig. S3C). Thus, increasing $\nu$ inevitably increases travel distance, and hence, the time to decision. Hence, for decisions where time to decision is critical, organisms require the spins in the decision-making ensemble to be highly tuned. However, very high directional tuning (low values of $\nu$) is also detrimental to the animal as this eliminates all options, except one, making the animal less sensitive to information that may be acquired as the animal approaches the targets. Thus, we expect the extent of neural tuning, together with neural noise, to represent a trade-off between decision-speed and accuracy.

**1.5 Calculating susceptibility.** Susceptibility is a key concept in physics and mathematics, from statistical mechanics to dynamical systems theory. It quantifies the change of a system's state in response to change in some external field. The reason physicists and mathematicians are interested in the quantity is because it exhibits a sharp increase when a complex system approaches a critical phase transition and is hence, predictive of the system being close to a tipping point (24, 25). For decision-making systems, susceptibility represents increased sensitivity of the system to small differences between the targets, making a rise in susceptibility highly desirable. Here, we show that the brain breaks down multi-choice decisions to a series of binary decisions, each of which is marked by a bifurcation, inevitably causing a peak in susceptibility. We measure susceptibility as the directional deviation of the agent given one extra spin that fires for one of the targets (making it $\sim 2\%$ more attractive). To do this we simulate a decision-making agent that at each timestep, outputs a desired direction of movement (following the exact procedure described above). Additionally, we introduce a second neural network composed of one extra spin encoding one of the targets, that at each timestep reaches equilibrium through iterative collective dynamics (here, for 1000 timesteps) and computes a second desired direction of movement. Susceptibility at the agent's current location is then measured as the difference in the agent's direction of travel output by the two networks (representing sensitivity to the presence of an extra spin in the second network). As expected, we observe a peak in susceptibility close to the bifurcation point. Quantifying the agent's decision accuracy reveals that the system is able to amplify such small differences and the agent almost always chooses the 'correct' target (Fig. S3, D and E).

**1.6 Spatial asymmetry in target locations.** In the above model formulation, spins encode target locations, and attractiveness of a target is encoded in the proportion of spins that point to that target. This formulation fails to encapsulate certain spatial geometries. For example, when two targets are in directional proximity while a third target is directionally distant from the agent's egocentric perspective. Since spins encode directions to targets, and two of the three targets are in directional proximity, the agent always goes to one of these targets (Fig. S8). Previous work in fruit flies (26), and our experiments with larval zebrafish (main Fig. 4), suggest that this is not the case and that animals integrate such that directionally-similar targets are considered as a single unit; that they will still visit an equally attractive, but directionally distant, target. To account for this, we incorporate an 'overlap' function in our model implementation. This function reduces effective size of the decision-making ensemble by discounting a proportion of spins encoding a target if there are other targets in directional proximity. Biologically, this can be seen as saturation of neurons that encode that direction. Here, each spin encodes direction to its preferred target with a Gaussian error, $\varepsilon_i$ centered around the direction to the target.

$$\varepsilon = \mathcal{N}(0, {\sigma_\theta}^2) \qquad [5]$$

where, ${\sigma_\theta}^2$ is the variance of the error distribution. Spins are then discounted from network depending on their deviation from their preferred target relative to their deviation from direction to other targets. Probability that spin $i$ is still considered, or probability that it is not discounted $c_i$ is given by

$$c_i = \frac{f(\theta_{ij})}{\sum_j f(\theta_{ij})} \quad [6]$$

where, $f(\theta_{ij})$ is the probability density at $\theta_{ij}$ from a normal distribution $N(0, {\sigma_\theta}^2)$ and $\theta_{ij}$ is the deviation in the preferred direction of spin $i$ from the center of target $j$. Effectively, a discounted spin in the decision-making ensemble and a spin that does not fire i.e. $\sigma_i = 0$ are treated identically. Fig. S4 shows results from a model with and without implementation of the 'overlap' function for symmetric setups (an illustration of including gaussian error in the directional vectors), while Fig. S8 shows these results in an asymmetric setup as discussed above.

**1.7 Mean-field approximation.** Here, we present a mean-field approximation of the same neural decision-making model described in section 1.2. This model largely draws inspiration from spin models used in physics, primarily to explain magnetism (1, 27). As analogy, neural activity here akin to spins in these models, excitatory neural interactions are described as being ferromagnetic and inhibitory interactions as antiferromagnetic.

In our model the $N$ spins are divided into $k$ equal groups $G_i (i = 1, \ldots, k)$, where $k$ is the number of options (potential targets in space) available to the animal. The fraction of the total number of spins that are active towards $\hat{p}_i$ is given by

$$n_i = \frac{1}{N} \sum_{j \in G_i} \sigma_j \quad [7]$$

Then we can rewrite equation [4] in the following way

$$\vec{V} = v_0 \sum_{i=1}^{k} \hat{p}_i n_i \quad [8]$$

The rule by which a spin switches its state from inactive ($\sigma_i = 0$) to active ($\sigma_i = 1$) is constructed such that the spin is more likely to be active if the animal is already moving in that direction. This can be expressed by Glauber dynamics (28).

$$r^{(i)}_{1\to0} = \frac{r_0}{1 + \exp\left(\frac{2k\vec{V}\cdot\hat{p}_i}{T}\right)} \qquad r^{(i)}_{0\to1} = \frac{r_0}{1 + \exp\left(-\frac{2k\vec{V}\cdot\hat{p}_i}{T}\right)} \quad [9]$$

where $r^{(i)}_{1\to0}$ is the rate in which a spin in group $G_i$ changes from "active" state to "inactive" and $r^{(i)}_{0\to1}$ is the rate of the opposite transition, from "inactive" to "active". $r_0$ is a constant rate which we set to one. The model also includes noise in the neural system i.e. the rate at which spins will switch states spontaneously independent of the collective dynamics involved. This is analogous to the temperature parameter that introduces randomness in the spin-flipping dynamics. Then the equations of motion (master equation) in the limit of $N \gg 1$ are

$$\frac{dn_i}{dt} = \frac{\frac{1}{k} - n_i}{1 + \exp\left(-\frac{2k\vec{V}\cdot\hat{p}_i}{T}\right)} - \frac{n_i}{1 + \exp\left(\frac{2k\vec{V}\cdot\hat{p}_i}{T}\right)} \quad [10]$$

We rearrange the above equation [10] to get

$$\frac{dn_i}{dt} = \frac{1}{k\left(1 + \exp\left(-\frac{2k\vec{V}\cdot\hat{p}_i}{T}\right)\right)} - n_i \quad [11]$$

The steady state solution of this equation can be written as the solution of the following system of algebraic equations

$$n_i = \frac{1}{k\left(1+\exp\left(-\frac{2k\vec{V}\cdot\hat{p}_i}{T}\right)\right)} \qquad i = 1, 2, ..., k \qquad [12]$$

The system of equations that including $k$ equations [12] and the 2 equations [8] in 2D is our basic system that gives us as its solution the velocity $\vec{V}$ and the fraction of active spins in each group $n_i$ at steady state. We will henceforth refer to this system as the "model equations".

When the targets are at infinity, the angles between the targets are constant, and the Hamiltonian is time-independent and describes a system in equilibrium. We now examine the simplest case of two targets at infinity, i.e. where $k = 2$ and $\hat{p}_1$ and $\hat{p}_2$ are fixed. In this case, there exists a symmetric solution that describes a compromise between the two targets.

$$\vec{V} = V\left(\frac{\hat{p}_1 + \hat{p}_2}{2}\right) \qquad [13]$$

Substituting equation [10] into equation [8] gives the following algebraic equation:

$$V = \frac{1}{1+\exp\left(-\frac{2V(1+\cos\theta)}{T}\right)} \qquad [14]$$

which always has a solution for $(0 < \theta < \pi;\ 0 < T < 1)$. In the three-choice case ($k = 3$), we get a similar compromise solution when the targets are radially symmetric, $\hat{p}_1 \cdot \hat{p}_2 = \hat{p}_3 \cdot \hat{p}_3$.

When the angle is large enough ($\theta > \theta_c$) and the temperature is low enough ($T < T_c$), there exists a second non-symmetric solution to the model equations that we term "decision" as it describes breaking the compromise between the targets and change in the direction of movement. The curve in the phase diagram where the second solution appears is called "binodal curve" or "coexistence curve" (represented by the dashed line; see Fig. 1B and E in main text). For larger angles, the symmetric solution becomes unstable. In the phase diagram this instability happens at the "spinodal curve" (represented by the solid line; see Fig. 1B and E in main text). Between the two curves we have the metastable region where both the compromise and decision are possible, and we expect a transition between them in a form of a bifurcation of the compromise solution. This is a typical hysteresis region of first order phase transition that at higher temperatures becomes second order. The transition to a second order phase transition happens at the tri-critical point where the coexistence and spinodal curves coincide into the second order phase transition curve (above the tri-critical point). In the case of three targets, we have at the first bifurcation point a phase transition between two different compromise solutions—for small angles we have a compromise between all three directions to the targets while for large angles we have a compromise between two of the three targets. Thus, the symmetry is broken sequentially.

Let us now find the criterion for instability of a general solution of the model equations. The dynamical equation for the velocity can be obtained using equations [8] and [11].

$$\frac{d\vec{V}}{dt} = \sum_{i=1}^{k}\frac{dn_i}{dt}\hat{p}_i = \sum_{i=1}^{k}\frac{1}{k\left(1+\exp\left(-\frac{2k\vec{V}\cdot\hat{p}_i}{T}\right)\right)}\hat{p}_i - \vec{V} \qquad [15]$$

Let $\vec{V} = \vec{V_0}$ be a solution of the model equations and consider a small perturbation to the velocity in the perpendicular direction to the velocity $\vec{V_0}$.

$$\vec{V} = \vec{V_0} + \epsilon\hat{n}^0 \qquad [16]$$

where, $\hat{n}^0$ is the normal to $\vec{V_0}$. Substituting into equation [15], expanding to first order in $\epsilon$ and taking the normal component, we get the following equation for the perturbation $\epsilon$.

$$\frac{d\epsilon}{dt} = -A\epsilon + \mathcal{O}(\epsilon^2) \quad [17]$$

where

$$A \equiv 1 - \frac{1}{2T}\sum_{i=1}^{k} \text{sech}^2\left(\frac{k\vec{V_0}\cdot\hat{p}_i}{T}\right)(\hat{n}^0\cdot\hat{p}_i)^2 \quad [18]$$

Therefore, the solution $\vec{V} = \vec{V_0}$ is stable if $A > 0$ and unstable if $A < 0$. Hence, the curve $A = 0$ for the compromise solution is the spinodal curve.

***1.7.1 Susceptibility.*** Susceptibility is defined as the response of the order parameter of a system to an external field (29). In this model, a bias towards one of the targets is equivalent to application of an external field in models of spin interactions (1). The order parameter in our model is the velocity component normal to the individual's previous direction of movement. Here, we examine its response to introduction of a small imbalance in the normal direction. Let us denote a small asymmetry in one of the neural groups as $\alpha$. Without loss of generality we take the first neural group $G_1$ (assuming $\hat{n}^0\cdot\hat{p}_i \neq 0$). Then the velocity can be written as

$$\vec{V} = \sum_{i=1}^{k} \frac{1}{k\left(1+\exp\left(-\frac{2k\vec{V}\cdot\hat{p}_i}{T}\right)\right)}\hat{p}_i + \alpha\hat{p}_1 \quad [19]$$

When we decompose the velocity to the initial velocity and the small perturbation $v_n$ in the normal direction $\hat{n}^0$

$$\vec{V} = \vec{V_0} + v_n\hat{n}^0 \quad [20]$$

in which case, the susceptibility $\chi$ (at constant temperature) is given by

$$\chi = \left.\frac{d(\vec{V}\cdot\hat{n}^0)}{d\alpha}\right|_{v_n=0,T=const.} \quad [21]$$

We substitute the decomposition equation [20] into the normal component of equation [19].

$$\vec{V}\cdot\hat{n}^0 = \frac{1}{k}\sum_{i=1}^{k}\frac{1}{1+\exp\left(\frac{-2k\vec{V_0}\cdot\hat{p}_i - 2kv_n(\hat{p}_i\cdot\hat{n}^0)}{T}\right)}\hat{p}_i\cdot\hat{n}^0 + \alpha\hat{p}_1\cdot\hat{n}^0 \quad [22]$$

Taking the derivative at $v_n = 0$ yields

$$d(\vec{V}\cdot\hat{n}^0) = \frac{1}{2T}\sum_{i=1}^{k}\text{sech}^2\left(\frac{k\vec{V_0}\cdot\hat{p}_i}{T}\right)(\hat{n}^0\cdot\hat{p}_i)^2 dv_n + (\hat{p}_1\cdot\hat{n}^0)d\alpha \quad [23]$$

Using the definition from equation [18] and the fact that $d(\vec{V}\cdot\hat{n}^0) = dv_n$, we can write the susceptibility as

$$\chi = \frac{\hat{p}_1\cdot\hat{n}^0}{A} \quad [24]$$

Thus, susceptibility diverges when $A \to 0$, i.e. on the spinodal curve (see Fig. S5). Based on this analysis, we arrive at the conclusion that maximal susceptibility is obtained when the bifurcation occurs at the highest possible angle—as late as possible from the perspective of an animal approaching the different spatial targets.

***1.7.2 Short-time response function.*** Let us consider that our system of $N$ spins is in the averaging/compromise regime. Since the order parameter of the system is the velocity component in the direction normal to the movement direction, we will denote it by $V_n$, so that a change in the direction of the movement will be a result of $V_n \neq 0$. At $t = 0$, we add a single spin that encodes direction to one of the targets, say $\hat{p}_1$. The short time response of the system is defined here as the response of the system as manifested in a change of the order parameter $V_n$ in a very short time, so short that it allows for only one spin to switch states. In other words we are looking for a change $\delta V_n$ over the firing time of a single spin $\delta t$ as a result of bias introduced by the additional spin. In doing this, we generalize a similar calculation that appeared for a one dimensional spin model in (27).

From equation [8], we see that the change in state of a single spin that encodes target $i$ (pointing towards $\hat{p}_i$ direction) makes the following change in the normal component of the velocity vector.

$$\Delta V_n = \frac{\hat{p}_i \cdot \hat{n}}{N} \tag{25}$$

First, we consider a simple limit $T \to \infty$ where there is no response. The probability of a spin in the group $i$ to be in the active state ($\sigma_i = 1$) is given by

$$P = \frac{r^{(i)}_{0\to 1}}{r^{(i)}_{0\to 1} + r^{(i)}_{1\to 0}} = \frac{1}{1 + \exp\left(-\frac{2k\vec{V}\cdot\hat{p}_i}{T}\right)} \tag{26}$$

which in the limit $T \to \infty$ gives $1/2$. We add a single spin that encodes direction to the target pointing towards $\hat{p}_1$. Therefore it contributes to $V_n$ according to equation [25]. In order to find the contribution of the rest of the spins in the decision-making ensemble, we have to multiply the probability that each spin is active (which is $1/2$ without response) by the number of spins in the group and the contribution of one active spin according to equation [25]. We therefore get

$$V_n = \sum_{i=1}^{k} \frac{1}{2} \cdot \frac{N}{k} \cdot \frac{\hat{p}_i \cdot \hat{n}}{N} + \frac{\hat{p}_1 \cdot \hat{n}}{N} = \frac{1}{2k} \sum_{i=1}^{k} (\hat{p}_i \cdot \hat{n}) + \frac{1}{N}(\hat{p}_1 \cdot \hat{n}) \tag{27}$$

When the targets are symmetric with respect to the direction of movement, the first term in equation [27] vanishes, and we get

$$V_n = \frac{\sin(\frac{\theta}{2})}{N} \tag{28}$$

where $\theta$ is the angle between the targets in the case of two targets, and the angle between the leftmost and the rightmost targets in the case of three targets.

Now let us return to the response of a single spin over time $\delta t$. At $t = 0$, we introduced an additional spin that encodes direction $\hat{p}_1$. Prior to any response, equation [27] gives the expression for $V_n$. We can write the contributions of the change in state of all spins in the ensemble in the following way which gives us the response of the system.

$$\delta V_n = \frac{1}{2} \cdot \frac{N}{k} \sum_{i=1}^{k} r^{(i)}_{0\to 1} \frac{\hat{p}_i \cdot \hat{n}}{N} \delta t - \frac{1}{2} \cdot \frac{N}{k} \sum_{i=1}^{k} r^{(i)}_{1\to 0} \frac{\hat{p}_i \cdot \hat{n}}{N} \delta t + \frac{\hat{p}_1 \cdot \hat{n}}{N} \delta t \tag{29}$$

Substituting the rates from equations [9], we get

$$\delta V_n = \sum_{i=1}^{k} \frac{\hat{p}_i \cdot \hat{n}}{2k} \tanh\left(\frac{k\vec{V}(0) \cdot \hat{p}_i}{T}\right) \delta t + \frac{\hat{p}_1 \cdot \hat{n}}{N} \delta t \tag{30}$$

Let us assume that we start from the symmetric compromise solution for $t < 0$, so that $V_n = 0$, and thus at $t = 0$, $V_n$ is still small. Then we can look at it as a small perturbation of $\vec{V}(0)$ according to equation [19], and obtain to first order in the perturbation $v_n$

$$\frac{\delta V_n}{\delta t} = \sum_{i=1}^{k} \frac{\hat{p}_i \cdot \hat{n}_0}{2k} \tanh\left(\frac{k\vec{V}_0 \cdot \hat{p}_i}{T}\right) + v_n(1 - A) + \frac{\hat{p}_1 \cdot \hat{n}_0}{N} \tag{31}$$

where $A$ is given in equation [18]. Since $A \geq 0$ for stable solutions, from the structure of the equation [31] we see that the response is maximal when $A = 0$, namely at the spinodal on the verge of instability (see Fig. S5).

Since there is no response yet at $t = 0$, we can substitute for $v_n$ the expression from equation [27]. Then for two and for three symmetric targets, we get

$$\frac{\delta V_n}{\delta t} = \frac{\sin\frac{\theta}{2}}{N}\left[1 + \frac{\sin^2\frac{\theta}{2}}{2T} \mathrm{sech}^2\left(\frac{2V_0 \cos\frac{\theta}{2}}{T}\right)\right] \tag{32}$$

and

$$\frac{\delta V_n}{\delta t} = \frac{\sin\frac{\theta}{2}}{N}\left[1 + \frac{\sin^2\frac{\theta}{2}}{T} \mathrm{sech}^2\left(\frac{3V_0 \cos\frac{\theta}{2}}{T}\right)\right] \tag{33}$$

respectively.

Now let us look at response due to the same bias that is introduced at $t = 0$, but $V_n(t < 0) \neq 0$ as it is essential in order to calculate the response function after the first bifurcation. In this case, $V_n$ is not small and there is a response for $t < 0$. We can add the additional spin that encodes direction $\hat{p}_1$ in the following way

$$\vec{V}(t = 0) = \vec{V}(t < 0) + \frac{\hat{p}_1 \cdot \hat{n}}{N} \hat{n} \tag{34}$$

where we denote $\vec{V}(t < 0) = \vec{V}_0$. We can write the response to this additional spin at $t = 0$ over the very short time $\delta t$ in the following schematic form

$$\delta V_n = \sum_{i=1}^{k} \frac{\hat{p}_i \cdot \hat{n}}{N} (P^{(1)}_{i,0\to1} - P^{(1)}_{i,1\to0}) \delta t + \frac{\hat{p}_1 \cdot \hat{n}}{N} \delta t \tag{35}$$

where $P^{(1)}_{i,0\to1}$ is the firing probability of an inactive spin in group $G_i$ as a response to bias introduced by the additional spin that encodes direction $\hat{p}_1$, and $P^{(1)}_{i,1\to0}$ is the probability that an active spin in group $G_i$ turns off as a response to bias introduced by the additional spin.

$$
\begin{aligned}
P^{(1)}_{i,0\to 1} &= \frac{N}{k} r^{(i)}_{1\to 0}(t<0) r^{(i)}_{0\to 1}(t=0) \\
P^{(1)}_{i,1\to 0} &= \frac{N}{k} r^{(i)}_{0\to 1}(t<0) r^{(i)}_{1\to 0}(t=0)
\end{aligned}
\qquad [36]
$$

where

$N/k$ is the number of spins in group $i$,

$P^{(i)}_{1\to 0}(t<0) = \frac{1}{1+\exp\left(\frac{2k\vec{V}_0\cdot\hat{p}_i}{T}\right)}$ is the probability of having an inactive spin at $t<0$ in group $G_i$,

$P^{(i)}_{0\to 1}(t<0) = \frac{1}{1+\exp\left(-\frac{2k\vec{V}_0\cdot\hat{p}_i}{T}\right)}$ is the probability of having an active spin at $t<0$ in group $G_i$,

$P^{(i)}_{1\to 0}(t=0) = \frac{1}{1+\exp\left(\frac{2k\vec{V}(t=0)\cdot\hat{p}_i}{T}\right)}$ is the probability of a spin to become inactive at $t=0$ in group $G_i$,

$P^{(i)}_{0\to 1}(t=0) = \frac{1}{1+\exp\left(-\frac{2k\vec{V}(t=0)\cdot\hat{p}_i}{T}\right)}$ is the probability of a spin to become active at $t=0$ in group $G_i$,

$P^{(i)}_{1\to 0}(t<0) r^{(i)}_{0\to 1}(t=0)$ is the probability of having an inactive spin at $t<0$ in group $G_i$, that becomes active at $t=0$,

$P^{(i)}_{0\to 1}(t<0) r^{(i)}_{1\to 0}(t=0)$ is the probability of having an active spin at $t<0$ in group $G_i$, that becomes inactive at $t=0$.

Substituting equation [34] into equation [35] and taking the limit of large $N (N \gg k)$, we get

$$
\begin{aligned}
\delta V_n = \sum_{i=1}^{k} \frac{\hat{p}_i\cdot\hat{n}}{N} \cdot \frac{N}{4k} \mathrm{sech}^2\left(\frac{k\vec{V}_0\cdot\hat{p}_i}{T}\right) \left[1 + \frac{2k(\hat{p}_1\cdot\hat{n})(\hat{p}_i\cdot\hat{n})}{NT\left(1+\exp\left(\frac{2k\vec{V}_0\cdot\hat{p}_i}{T}\right)\right)}\right]\delta t \\
- \frac{\hat{p}_i\cdot\hat{n}}{N} \cdot \frac{N}{4k} \mathrm{sech}^2\left(\frac{k\vec{V}_0\cdot\hat{p}_i}{T}\right) \left[1 - \frac{2k\exp\left(\frac{2k\vec{V}_0\cdot\hat{p}_i}{T}\right)(\hat{p}_1\cdot\hat{n})(\hat{p}_i\cdot\hat{n})}{NT\left(1+\exp\left(\frac{2k\vec{V}_0\cdot\hat{p}_i}{T}\right)\right)}\right]\delta t + \frac{\hat{p}_1\cdot\hat{n}}{N}\delta t \\
= \frac{\hat{p}_1\cdot\hat{n}}{N}(2-A)\delta t
\end{aligned}
\qquad [37]
$$

where $A$ is given by equation [18]. Also in this case we see that the response is maximal when $A=0$, namely at the spinodal on the verge of instability (see Fig. S5).

***1.7.3 Trajectories.*** We use the same model equations for a time-dependent Hamiltonian form [1] and consider them at every position to calculate trajectories for a system with targets at a finite distance (see Fig. S5 for trajectories obtained from the mean-field approximation and Fig. S6 for comparison with simulations of smaller system sizes).

All results of this section can be generalized to any value of the neural tuning parameter by replacing the cosines of the scalar products between the directions $\hat{p}_i$ by the expression given in equation [2].

**1.8 Embedding the neural model in the ring-attractor class of models.** Much work in neuroscience has considered ring-attractor models as being of considerable importance to understanding spatial navigation (4, 5, 16, 30–32). Neural ring-attractor models (4, 31) are motivated by head-direction cells found in the brain of both vertebrates (23, 33) and invertebrates (16, 30). They are known to underlie the representation of instantaneous heading direction of an animal in the horizontal plane regardless of its location and ongoing behavior (34). When animals are exposed to a prominent landmark, an activity bump appears on a specified sector of the ring, and rotates concurrently with the landmark as the animal turns towards it (30). Upon introduction of a second landmark, this activity bump was found to either 'flow' or 'jump' to, and stabilize at, a new location. The exact nature of the bump shift ('flow' or 'jump') was found to be dependent on the angular distance of the new landmark from the older one (16). All these dynamics are captured by neural ring-attractor models. We follow the general structure of the neural ring-attractor models presented for example in (5) and (32).

In the ring-attractor model, neural firing rate is described in a continuous formulation by a single scalar function $m(\theta, t)$ which is a function of the preferred angle $\theta$ and time $t$. The model is essentially one-dimensional with ring topology, and the distribution of the preferred directions is treated as continuous and uniform. The ring is defined by the preferred directions and thus proximity of neurons in the ring does not imply physical proximity in the brain, although we note that in some insects, directions are topologically organised (16, 30). The interactions in the network are such that an activity bump is created and maintained by excitation among neurons that encode a similar direction, along with global inhibition or inhibition among neurons that encode conflicting directions (35). Since our model inherently includes excitation when the spins encode a similar direction and inhibition in different directions, it becomes natural to identify the (average) firing rate with the fraction of active spins in the mean field limit, and below we show that our model is within the class of neural ring-attractor models.

The dynamics of the firing rate is expressed schematically by the following equation

$$\tau \frac{\partial m(\theta,t)}{\partial t} = -m(\theta,t) + g\left(I(\theta,t) + X(\theta,t)\right) \quad [38]$$

where $g$ is a non-linear function which describes the gain of a spin from the input which can further be split into interactions with rest of the network $I(\theta,t)$ and external input $X(\theta,t)$ and $\tau$ is a time constant that sets the time scale of the dynamics of the system (32). The gain function $g$ is usually taken to be Heaviside step function (5, 32).

Interactions with the rest of the network are usually given as a convolution of the following form

$$I(\theta,t) = \frac{1}{\pi}\int d\theta' J(\theta-\theta')p(\theta',t)m(\theta',t) \quad [39]$$

where the function $J$ describes the relative weights of the neurons in the network. It commonly takes the form of a Ricker wavelet, also known as a Mexican Hat function, which has an excitatory center and an inhibitory surround, usually depend only on the difference in orientation. $p(\theta,t)$ denotes the release probability of a neuron which represents the efficiency of the neural interactions.

Below, we illustrate how our neural model suggests a mixture between the two input terms (the interactions and the external input). Since the gain function acts on both, there is no concrete linearity that should be preserved. Mathematical formulations of the ring-attractor network sometimes include a stochastic function in the form of a $\theta$-dependent Wiener process. This is added to [38] to represent external noise (36). The neural model gives a natural way to extend this model to include noise presented in our model as the neural noise parameter $T$.

Let us consider $N$ spins at each point of the ring. In this regard we take a continuous limit of the neural model, and in the end we consider the mean field limit, which means in particular taking $N \gg 1$.

$$H = -\frac{J}{2N}\int d\theta' \int d\theta \sum_{i=1}^{N}\sum_{j=1}^{N} \sigma_i(\theta,t)\sigma_j(\theta',t)\hat{p}_i\cdot\hat{p}_j + \eta\int d\theta \sum_{i=1}^{N}\sigma_i(\theta,t) - \eta\sum_{l=1}^{k}\int d\theta \sum_{i=1}^{N}\delta(\theta-\bar{\theta}_l)\sigma_i(\theta,t) \quad [40]$$

$J$ is the interaction strength which is assumed to be equal for all spins in all to all interactions. The interactions here (first term in [40]) are considered to be as the regular scalar products between different directions $(\hat{p}_i(\theta,t)\cdot\hat{p}_j(\theta',t))$ which can be modified according to equation [2] to include the neural tuning parameter $\nu$. The parameter $\eta$ represents global inhibition (second term in [3]) that takes effect at any angle $\theta$ and it is large enough to prevent the spins from changing their state from 0 to 1 ($\eta N \gg J$), unless an external stimuli induces an effective external field that cancels out the effect of the global inhibition at those angles (third term in [40]). See (37) for a similar construction with inhibition and induced external field for a neural network. The stimuli represent the targets that positioned at the angles $\bar{\theta}_l(t)$. We assume that they induce a very narrow distribution of external field centered at $\theta = \bar{\theta}_l$ $(l = 1,..,k)$, that we write them here with delta functions.

In order to obtain the free energy in the mean field limit we can rewrite the Hamiltonian in the following way

$$H = -\frac{J}{2N}\int d\theta' \int d\theta \left(\sum_{i=1}^{N}\sigma_i(\theta,t)\hat{p}_i\right)\cdot\left(\sum_{j=1}^{N}\sigma_j(\theta',t)\hat{p}_j\right) + \eta\int d\theta \sum_{i=1}^{N}\sigma_i(\theta,t) - \eta\sum_{l=1}^{k}\int d\theta \sum_{i=1}^{N}\delta(\theta-\bar{\theta}_l)\sigma_i(\theta,t) \quad [41]$$

We are following here the procedure that was used for the one dimensional Curie-Weiss model (see (38), chapter 13) and was applied in the context of a similar spin model Hamiltonian in (1) for the case of two targets. The partition function for the Hamiltonian is

$$Z = tr_{\{\sigma_i\}}\exp\left(-\frac{H}{T}\right) \quad [42]$$

Since the system is two-dimensional, let us introduce two auxiliary fields $\vec{V} \equiv (V_x, V_y)$ and using the Gaussian identity

$$e^{\frac{a^2+b^2}{4\,c}} = \frac{c}{\pi}\int_{-\infty}^{\infty} dV_x dV_y\, e^{-c\,(V_x^2+V_y^2)+a\,V_x+b\,V_y} \qquad [43]$$

where we take

$$\begin{aligned} a &= \frac{2\,J^{\frac{1}{2}}}{T}\int d\theta \sum_{i=1}^{N}\sigma_i(\theta,t)\,\cos\theta, \\ b &= \frac{2\,J^{\frac{1}{2}}}{T}\int d\theta \sum_{i=1}^{N}\sigma_i(\theta,t)\,\sin\theta, \\ c &= \frac{N}{T} \end{aligned} \qquad [44]$$

We can write the partition function in a form which is linear in $\sigma_i$

$$Z = \frac{N}{\pi\,T}\,tr_{\{\sigma_i\}}\int_{-\infty}^{\infty} dV_x dV_y \exp\left(-\frac{N\,\vec{V}^2}{T} + \frac{1}{T}\int d\theta\left(2J^{\frac{1}{2}}\sum_{i=1}^{N}\sigma_i\,\widehat{p}_i(\theta)\cdot\vec{V} - \eta\left(1-\sum_{l=1}^{k}\delta\left(\theta-\bar{\theta}_l\right)\right)\sum_{i=1}^{N}\sigma_i\right)\right) \qquad [45]$$

In the limit of large $N$ the partition function is dominated by

$$\vec{V} = J^{\frac{1}{2}}\int d\theta n(\theta)\widehat{p}(\theta) \qquad [46]$$

where $n(\theta)$ is the fraction of active spins defined by $n(\theta) \equiv \frac{1}{N}\sum_{i=1}^{N}\sigma_i(\theta)$. Summing over the possible states of the spins we get

$$\begin{aligned} Z &= \frac{N}{\pi\,T}\int_{-\infty}^{\infty} dV_x dV_y e^{-\frac{N\,\vec{V}^2}{T}}\int \mathcal{D}\theta\prod_{i=1}^{N}\sum_{\sigma=0,1} exp\left(\frac{1}{T}\left[2\,J^{\frac{1}{2}}\,\vec{V}\cdot\widehat{p}_i(\theta) - \eta(1-\sum_{l=1}^{k}\delta(\theta-\bar{\theta}_l))\right]\sigma\right) \\ &= \frac{N}{\pi\,T}\int_{-\infty}^{\infty} dV_x dV_y e^{-\frac{N\,\vec{V}^2}{T}} e^{N\sum_\theta \ln\left(1+\exp\left[\frac{1}{T}\left(2\,J^{\frac{1}{2}}\,\vec{V}\cdot\widehat{p}(\theta)-\eta\left[1-\sum_{l=1}^{k}\delta_{\theta\bar{\theta}_l}\right]\right)\right]\right)} \end{aligned} \qquad [47]$$

where $\mathcal{D}\theta \equiv \prod_\alpha d\theta_\alpha$ over all angles.

Then we can read the free energy per spin $F(\vec{V},T)$ from the general form

$$Z \sim \int_{-\infty}^{\infty} dV_x dV_y \exp\left(-\frac{N}{T}F(\vec{V},T)\right) \qquad [48]$$

to be

$$F(\vec{V},T) = \vec{V}^2 - T\sum_\theta \ln\left(1+\exp\left[\frac{1}{T}\left(2J^{\frac{1}{2}}\,\vec{V}\cdot\widehat{p}(\theta) - \eta\left[1-\sum_{l=1}^{k}\delta_{\theta\bar{\theta}_l}\right]\right)\right]\right) \qquad [49]$$

The effective result of this construction is that at each value of $\theta$ along the ring we obtain a separate contribution to the equation of motion in the mean field approximation. From the equations of motion $\frac{\partial F}{\partial \vec{V}} = 0$ we get the steady state solution

$$\vec{V} = \sum_{\theta} \frac{J^{\frac{1}{2}}\widehat{p}(\theta)}{1 + \exp\left(-\frac{1}{T}\left[2\,J^{\frac{1}{2}}\vec{V}\cdot\widehat{p}(\theta) - \eta\left(1 - \sum_{l=1}^{k}\delta_{\theta\bar{\theta}_l}\right)\right]\right)} \qquad [50]$$

and according to equation [46] we identify the fraction of active spins at steady state to be

$$n(\theta, t) = \frac{1}{1 + \exp\left(-\frac{1}{T}\left[2J\int d\theta' n(\theta', t)\widehat{p}(\theta', t)\cdot\widehat{p}(\theta, t) - \eta\left(1 - \sum_{l=1}^{k}\delta_{\theta\bar{\theta}_l}\right)\right]\right)} \qquad [51]$$

We see here that indeed if the global inhibition is high, $n(\theta, t) \to 0$, except for $\theta = \bar{\theta}_l$ $(l = 1, .., k)$. The corresponding dynamical equation in analogy with equation [11] is

$$\frac{\partial n(\theta, t)}{\partial t} = \frac{1}{1 + \exp\left(-\frac{1}{T}\left[2J\int d\theta' n(\theta', t)\widehat{p}(\theta', t)\cdot\widehat{p}(\theta, t) - \eta\left(1 - \sum_{l=1}^{k}\delta_{\theta\bar{\theta}_l}\right)\right]\right)} - n(\theta, t) \qquad [52]$$

and the resemblance in structure to equation [38] leads us to the identification of the fraction of active spins $n(\theta, t)$ with the firing rate in the ring model. Thus, our model is within the class of neural ring-attractor models. However, it has the advantage of being simple and analytically tractable. Furthermore, much experience that comes from analyzing such spin systems has lead to their wide adoption in neuroscience (20, 22, 39).

### 2. Collective decision-making model for animal groups

**2.1 Overview.** The bifurcation pattern described above for binary decision-making is reminiscent of work on collective decision-making in bird flocks (40), fish schools (41) and baboon troops (42). We expose an established model of consensus decision-making in animal groups (43) to a multi-choice decision scenario. In the absence of any feedback mechanism, we find that this model will fail to produce the bifurcation patterns observed in our data. However, previous work has shown that uninformed individuals (those without a desired direction of travel), to some degree, are able to provide this feedback, at least when the group is deciding between two options (43)). However, because the uninformed individuals are recruitable by informed individuals with different desired directions of travel, they primarily function to keep the group together, and fail to provide the necessary feedback in the three-choice context (Fig. S19. Thus, the group will almost always approach the central target. By introducing feedback on individual goal-orientedness as a function of their experienced travel direction, we are able to produce all bifurcation patterns observed in our experimental system, in the presence of two, and three, options. With these feedback mechanisms in place, the model predicts that animal groups, like the brain, will break multi-choice decisions to a series of binary decisions (see Fig. 5 in main text). Thus, our results may be broadly applicable across scales of biological organization—neural ensembles and animal collectives. At both these scales, decision-making can be conceived as a consensus paradigm among elements that compose the system—consensus among neurons at the individual level and among individuals at the group level.

**2.2 Framework.** Groups are composed of $N$ individuals, each characterised by a position vector $c_i(t)$ and unit direction vector $\hat{v}_i(t)$ where $i$ is the individual identity and $t$ is the current time step. Individuals modify their own motion by responding to neighbours within a certain distance from them. They turn away from $n_r$ neighbours encountered within a small repulsion zone of radius $r_r$. This represents collision avoidance and maintenance of personal space, and as is apparent in real animal groups, takes highest priority.

$$\vec{d_i}(t + \Delta t) = -\sum_{i=1, i\neq j}^{n_r} \frac{c_j(t) - c_i(t)}{|c_j(t) - c_i(t)|} \qquad [53]$$

where $d_i(t + \Delta t)$ represents the individual's desired direction of travel in response to conspecifics. If no neighbor is present in this zone, the focal individual is attracted to and aligns with $n_a$ neighbours within a larger interaction zone of radius $r_a$.

$$\vec{d_i}(t + \Delta t) = \sum_{i=1, i\neq j}^{n_a} \frac{c_j(t) - c_i(t)}{|c_j(t) - c_i(t)|} + \sum_{j=1}^{n_a} \frac{v_j(t)}{|v_j(t)|} \qquad [54]$$

Here, $d_i(t + \Delta t)$ is subsequently converted to the corresponding unit vector $\hat{d}_i(t + \Delta t) = d_i(t + \Delta t)/|d_i(t + \Delta t)|$. To incorporate target preferences, individuals are given information about a preferred direction. Each individual is attributed a goal vector $g_i(t)$ that points to one of the targets amongst which the group must choose. For sake of simplicity, we assume all individuals have a preferred target, and that the number of individuals with preference for a given target is the same as the number of individuals with preference for any other target. Individuals balance this personal preference with social interactions using a weighting term $\omega$ to give their desired direction of travel.

$$\vec{d_i}'(t + \Delta t) = \frac{\hat{d}_i(t + \Delta t) + \omega g_i(t)}{|\hat{d}_i(t + \Delta t) + \omega g_i(t)|} \qquad [55]$$

Motion of all individuals is subject to noise (error in movement and/or sensory integration) which is implemented by rotating $\vec{d_i}'(t + \Delta t)$ by a random angle chosen from a circularly wrapped Gaussian distribution centered at 0 and of standard deviation $\sigma_e$. Once the desired direction is determined, individuals turn towards $\vec{d_i}'(t + \Delta t)$ with a maximum turning rate of $\psi \Delta t$.

As in (43), we implement feedback on $\omega$. At each timestep, if individuals find themselves moving in the direction of their preferred motion (here, within 20° of their preferred direction), $\omega$ is reinforced by a small value $\omega_{inc}$ until it reaches a maximum value $\omega_{max}$. Otherwise, it is reduced by $\omega_{dec}$ until it reaches 0. Since the group stays together, group movement towards a given option activates individuals with similar directional preferences by increasing their $\omega$ while inhibiting individuals with opposing directional preferences by decreasing their $\omega$.

We also perform simulations without feedback on $\omega$ to emphasise that this is an essential feature for the model to produce bifurcations based on egocentric geometry of the presented options (Fig. S19). Without the feedback, the model predicts that animal groups will rarely leave the averaging regime and that groups tend to split. For the three-choice case, in the few cases where the group does not split, it approaches the middle target (Fig. S19).

**2.3 Simulations.** We simulated a group of sixty agents exposed to two or three targets in their environment. Individuals were initialized in proximity to (0, 0) with random positions and directions. For the two-choice case, targets were positioned at a distance of 5 units and 60° apart from the group's perspective (corresponding to target locations in the neural model, and the fly experiments). This places the two targets at locations (4.33, -2.5) and (4.33, 2.5) respectively. For three targets, distance was still maintained at 5 units but successive targets were now placed 40° apart. The targets were now located at (3.83, -3.21), (5, 0) and (3.83, 3.21). All individuals were assigned a preferred target randomly such that each target had equal number of individuals whose goal vector pointed to it. We ran simulations with and without feedback on the $\omega$ term to show that this feature is essential to produce such patterns (Fig. S19). We ran 500 replicate simulations for each condition and each replicate run was considered successful when the group reached a given target without splitting. In order to minimize group splits, these simulations were run with 12 informed individuals having preference each target. The remaining individuals (36 and 24 individuals for the two- and three-choice simulations respectively) were considered 'uninformed' and exhibited no preference to any target.

### 3. Experiments with fruit flies (*Drosophila melanogaster*)

**3.1 Fly preparation.** All experiments were conducted on 3- to 5-day old female wild-type CS strain *Drosophila melanogaster* raised at 26 °C on a 12 hr light, 12 hr dark cycle. Prior to the experiment, individual flies were anaesthetized in an icebox. The anaesthetized flies were placed on a Peltier stage maintained at 4 °C and glued to a 0.15 mm V2A stainless steel pin using UV-curing glue. The animals were given at least 20 min to recover from the anesthesia before introducing them to the experimental setup. All experiments were carried out in the last 4 hr of the animal's subjective day at 20 °C.

**3.2 Fly virtual reality experiments.** Experiments were conducted in a flyVR setup procured from loopbio GmbH. Tethered flies were positioned in the center of an acrylic bowl of diameter 20 cm, lowered 7 cm from the bowl surface. The bowl is used as a hemispherical projection surface for the visual stimuli. Flies were filmed from an angle using a camera (Basler acA645-100gm; lens: kowa 25 mm/f 1.4) equipped with an infrared filter (Lee filters, transmission above 730 nm) at 100 Hz and were illuminated using infrared light at 850 nm to track their heading direction. We assume that the fly flies at constant speed of 0.2 m/s in this direction to close the loop in our 2D virtual reality setup. A constant speed assumption causes the visual stimulus to update even when the fly stops flapping its wings. To ensure that the fly flies (flaps its wings) during the entire course of the experiment, the experimenter gently blew on it when it stopped flying. Trials where flies stopped flying more than five times during the course of the experiment were excluded from further analyses. This step in data filtering distinguishes our fly data from the locust and fish data, where the VR systems are designed for freely-moving animals. It results in us selecting for flies that are inclined to fly (flap their wings) consistently, causing the raw trajectories to have lesser noise from a stop and go motion. However, it is also worth noting that this filtering still chose all individuals as long as they flew, regardless of whether or not they flew to a target.

Other reasons for a fly to be excluded from analyses were: i) Too much glue on the tether prevented the fly from flapping its wings freely, making it unsuitable for the experiment. ii) Too little glue on the tether would occasionally result in flies freeing themselves and flying away. iii) While our experimental room was temperature controlled, humidity varied over the course of the experiment. On days of low humidity, an external humidifier was used. Thus, conditions were not always ideal for a fruit-fly to "want" to flap its wings and fly. iv) The fly did not perform the stripe fixation task.

**3.3 Visual stimuli.** Custom 3D scenes were designed using 3D modelling program Blender (version 2.77) and projected on the bowl with a projector (Optoma ML750e DLP) at 120 Hz refresh rate. The stimulus created was a white 'shadeless' cube of side 50 m that served as background and ten black cylinders of 1m diameter each. Making the objects shadeless removes interaction of the object with light and hence removes any edges that may otherwise be visible on the cube. The position of each cylinder was determined from an SQLite database that was generated automatically (see experimental design for details). Pillars that were not part of the current stimulus were placed at >100 m distance where they were visually occluded by the cube.

**3.4 Data collection.** Tethered *Drosophila melanogaster* were exposed to either a two-choice or a three-choice decision task in the virtual reality environment. Each experimental trial lasted 15 min where flies were exposed to five stimuli—three experimental stimuli and two control stimuli. The experimental stimuli consisted of two or three cylinders (depending on the experimental condition) that were presented to the animal in three different angular conditions. The order in which the stimuli were presented were randomized. The control stimulus was presented before and after the experimental conditions. This was a stripe fixation task where the fly was exposed to a single cylinder and was expected to orient and fly towards this cylinder. This is a well-known response in tethered Drosophila and flies that did not perform this were excluded from further analyses. The actual position of cylinders in all stimuli were randomized rotationally to prevent effects of any directional bias that may arise from the geometry of the physical setup around the fly. In all conditions, the position of the fly was reset to the origin once it reached a cylinder or flew a corresponding distance in any direction. To ensure true randomizations during our experiment, we implemented them prior to starting any experimentation. An SQLite database was created which had positions of all posts for all experiments. A total of 60 flies were tested in the VR setup. Of these, 30 flies were exposed to a stimulus that consisted two targets, and 30 flies were exposed to a stimulus that consisted three targets. Our analyses include 70% and 74% of all the tracks adopted by flies in the two-choice and three-choice decision-making scenarios respectively. The remaining tracks were excluded because these were trials where none of the targets were approached by the fly within the remaining experimental duration, and hence, were considered to not constitute a decision-making scenario.

**3.5 Data analysis.** Each trajectory of a fly (travelled in virtual space from the origin till the position when the location of the fly was reset) was considered to be an event. We rotated all trajectories such that the x-axis points from the origin, to the center of mass of the targets (see Fig. S11 for fly trajectories in the presence of two and three options). To visualize trajectories in the various experimental conditions, we create time-normalized (proportion of maximum) density plots. Each density plot was constructed by picking, for each pixel, the maximum value among normalized density plots (proportion of maximum) for varying times in a sliding time window. In the two-choice case, flies show no bias towards either target, and the trial results are symmetric about the $x-$axis. For the three-choice case, we observe some asymmetry that may arise from random perturbations that inherently affect individual trajectories. We symmetrize our data by mirroring it about $y = 0$ to remove this asymmetry (based on the two-choice results). Next, to quantify the decision points, we fold the data about the line of symmetry, $y = 0$. We then applied a density threshold to the time-normalized (proportion of maximum across a sliding time window) density plot to reduce noise and fit a piecewise phase transition function to quantify the bifurcation.

$$y = \begin{cases} 0 & x \le x_c \\ A|x - x_c|^{\alpha} & x > x_c \end{cases} \quad [56]$$

where $x_c$ is the critical bifurcation point, $\alpha$ is the critical exponent, and $A$ is the proportionality constant. To avoid bias in the fit that arises from $y = 0$ part of the data (to the left of the bifurcation), we exclusively fit the above function in a range starting near to the suspected bifurcation point. For the three-choice case, the piecewise function is fit to each bifurcation separately. Additionally, for each bifurcation we also performed randomization tests where we repeated the exact fit procedure described above to data where the trajectories were randomized by keeping the $x-$coordinates, and swapping the $y-$coordinates with values from other random events. Occurrences of a bifurcation is then assessed using the following criteria: (a) The bifurcation occurred between $x = 0$ and $x = T_x$, where $T_x$ is the $x-$cooridnate of the two targets in consideration, (b) The critical exponent $0.2 < \alpha < 2$, and (c) The proportionality constant $A > 0.2$. Based on these criteria, randomizations showed that the resultant fit to our experimental data were highly significant ($p < 0.01$ for binary choice and $p < 10^{-4}$ for the three-choice case).

Based on the amount of time it took a fly to reach one of the available targets, we also classified individual fly tracks into one of two categories—direct tracks and non-direct tracks. This allowed us to quantify the proportion of tracks where the individuals exhibited the bifurcations predicted by our model (see Fig. S11, A and H). Such between-individual variability in response is expected, and it is very difficult to ascertain its source. Recent work with fruit flies (44, 45) has shown non-heritable variation in neural connectivity results in behavioral differences in individual tendency to orient, and move, towards a vertical stripe target. In our model, such differences can be accounted for by variation in the directional tuning of spins. A high neural tuning (low $\nu$) results in more directed tracks (Fig. S14).

### 4. Experiments with desert locusts (*Schistocerca gregaria*)

**4.1 Locust preparation.** All experiments were conducted on instar 5 desert locusts (*Schistocerca gregaria*) raised in the Animal Research Facility of the University of Konstanz. Locusts were moved to the experimental room one night prior to the experiment, where they were maintained at 26 °C. Experiments were then conducted at ∼31 °C and $20 - 22\%$ relative humidity with fully intact locusts. Each locust was used only once.

**4.2 Locust virtual reality experiments.** Experiments were conducted in a locustVR setup procured from loopbio GmbH (Fig. S10). The setup consists of three main components: (a) the locomotion compensator, (b) the recording system with a closed-loop extension, and (c) the FreemoVR system (46).

The locomotion compensator (a) is a two-dimensional treadmill composed of a hollow polyethylene sphere of diameter 60 cm. Two servo-motors with rotary encoders turn the sphere to compensate for the animal's movement, allowing it to move infinitely on the sphere (mechanics are adapted from (47)). The recording system with closed-loop extension (b) consists of a recording unit—a 100 fps infrared machine vision camera and an LED spotlight at 850 nm (infrared) and functions as the tracking and feedback-loop component of the system. Tracking the animal's movement facilitates feedback to the locomotion compensator to keep the animal centered on the sphere. Optical tracking is performed using a contrast-based method. The optical center of mass of the animal is detected and its deviation from the center of the sphere is converted into a compensation response of the motors. Finally, the closed-loop extension software feeds the animal movement to the VR system (c) to update projection of the stimulus accordingly. A vertical cylinder (70 cm tall; 80 cm diameter) was used as the projection surface and projections were done using three Optoma GT1070Xe projectors with overlapping projections. From the animal's perspective, this projection surface covers 360° field-of-view horizontally, and 74.9° vertically.

**4.3 Visual stimuli.** Custom 3D scenes were designed using 3D modelling program Blender (version 2.77) and projected on the projection surface. The stimulus created was identical to what is described above for flies (see Section 3.3), with the exception that the target cylinders for the locusts were of diameter 0.2 m.

**4.4 Data collection.** The data collection procedure for the desert locusts was identical to the procedure adopted for flies (see Section 3.4) except each experimental trial lasted 48 min—the three experimental stimuli lasted 12 min each, and the two control stimuli lasted 6 min each. As with the flies, the control stimulus was a stripe fixation task for the two-choice experiments. For the three-choice experiments, however, this was modified to be a two-choice decision task. A total of 156 locusts were tested in the VR setup. Of these, 57 locusts were exposed to stimulus that consisted two targets, and 99 locusts were exposed to a stimulus that consisted three targets (see Fig. S12 for locust trajectories during decision-making in the presence of two and three options). Based on our data filtering criteria, 122 out of the 156 locusts were used in the analyses. Of these, 35 locusts performed decision-making in the presence of two targets and 87 locusts performed decision-making in the presence of three targets. Of the locusts that were included in our analyses, 31% and 29% of tracks were used for further analyses in the two-choice and three-choice decision-making scenarios respectively. Note that this number is much smaller than it was for flies (70% and 74% for two- and three-choice experiments respectively). This difference arises from the additional data filtering step for the flies. Because the flyVR requires the animal to be tethered and assumes that it flies at a constant speed in the direction where it is facing, we exclude any animal that stops flying (flapping its wings) more than five times. The locustVR however is designed for freely walking animals. Hence, the animal is free to stop, and not move towards any of the targets presented to them. Because, the animal did not choose any of the presented targets, we consider these tracks to not constitute a decision-making scenario.

**4.5 Data analysis.** The analyses procedure adopted for the desert locusts was identical to the procedure adopted for flies (see Section 3.5). Randomizations performed on locust trajectories also showed that the resultant fit to these experimental data were highly significant ($p < 0.01$ for binary choice and $p < 10^{-4}$ for the three-choice case).

Similar to the flies, the locust trajectories were also classified as direct, or non-direct tracks. However, because the locustVR system allowed the animals to stop and reconsider movement during the decision-making process, we added an additional category to classification of individual locust tracks viz. the wandering tracks (Fig. S12, A and J). Once again, we find that variability in directedness of locust tracks predominantly occurs at the individual level. This type of non-heritable behavioral variability in space-use is expected, and is shown to be the case in many species including flies (44, 45), fish (48), pea aphids (49), mice (50) and crayfish (51).

### 5. Experiments with larval zebrafish (*Danio rerio*)

**5.1 Fish preparation.** All experiments were conducted on 1 cm ± 0.1 cm long zebrafish (*Danio rerio*) of age 24 to 26 days post-fertilization raised in a room at 28 °C on a 16 hr light, 8 hr dark cycle. The fish were bred and raised by the animal care staff of the Department of Collective Behaviour at the Max Planck Institute of Animal Behavior and the University of Konstanz. Fish were transferred to the experimental room at least 12 hr prior to the experiments in water from their holding tanks. This ensured that the water quality and temperature in the experimental room was the same as in their holding facility. This water was also used in the VR setups where a water change was done once a day. All fish were tested individually. They were naive, and chosen at random from their holding tanks. All experiments were conducted in accordance with the animal ethics permit approved by Regierungsprasidium Freiburg, G-17/170.

**5.2 Fish virtual reality experiments.** Experiments were conducted in a fishVR setup procured from loopbio GmbH (See (46) for setup details). Larval zebrafish were tested in an acrylic bowl of diameter 34 cm between 07:00 and 19:00. Once a fish was introduced in the arena, it was given 20 min to acclimatize to the environment. This was followed by a 10 min control where it was presented a single virtual conspecific circling the arena in a circle of radius 8 cm (Fig. S16). The purpose of this control was to assess whether the real fish would follow a virtual conspecific. We later included data from all fish in our analysis as nearly all of them followed the virtual fish during this control. After the control, the real fish was exposed to choice experiments that lasted 1.5 hr with the virtual fish initialized with random lateral distances between them and random swim direction.

**5.3 Visual stimuli.** Custom 3D scenes were designed using 3D modelling program Blender (version 2.77) and projected on the bowl with a projector (Optoma ML500) at 100 Hz refresh rate. The stimulus created was a larval zebrafish of length 1 cm. The fish was set to swim at an average speed of 4 cm/s with burst-and-glide motion extracted from a random real fish's swimming pattern (Fig. S16). The background for the stimulus was light blue, the default projection color on the Optoma projector.

**5.4 Data collection.** As choice experiments, the real fish was exposed to two or three virtual conspecifics that swam side-by-side with lateral distance between them varying from 0.5 cm to 10 cm (in steps of 0.5 cm). The virtual fish swam back-and-forth at a 3 cm depth, and along a straight line of length 24 cm (Fig. S15). To exclude boundary effects and the effect of sharp turns that the virtual fish make near the edge, we only consider data where the virtual fish are farther than 5 cm from the boundary for further analyses. The following filters were used to extract data where the real fish was considered to be potentially interacting with virtual conspecifics:

1. A distance filter determined whether the real fish had the opportunity to interact with a virtual fish. If the real fish was within 5 cm front-back distance or 5 cm left-right distance of the outermost virtual fish, it was considered to be within interaction range.

2. Since the real fish receives no feedback from its virtual conspecifics, we only consider cases where the virtual fish lead the real fish i.e. cases where the virtual fish are ahead of the real fish in a coordinate system centered at the real fish's frame of reference, and where the real fish is behind the virtual fish in a coordinate system centered at the centroid of the virtual fish and pointing in the direction of their motion.

3. Since we are interested in which conspecific(s) the real fish will follow, we exclude all data where the angle between the real fish's direction and the virtual fish's direction ($\phi$) is larger than 30°.

4. Finally, for the purpose of analysis, we switch identities of the virtual fish after each turn. This is done to ensure relative positions of the virtual fish are conserved; that virtual fish are always treated as being to the left or right of the real fish.

A total of 440 fish were tested. Of these, 198 fish were exposed to decision-making with two virtual targets, 39 fish were exposed to decision-making with three targets, and 50 fish were exposed to decision-making with three targets in asymmetric geometry. In the two-choice case, the real fish experienced five different virtual fish speeds. Our analyses focus only on data where the virtual fish swim at an average speed of 4 cm/s, the average swim speed of larval zebrafish. For the experiments in asymmetric geometry, the real fish was exposed to choice experiments where distance between the center virtual fish and its closer neighbor was 0.03 m and its distance to the other neighbor was 0.09 m (see main Fig. 4). Note that the left-right position of the two fish closer to each other was randomized, indicating that this pattern did not result from handedness in individual fish. We also conducted experiments where the real fish was exposed to a single virtual fish, or where two real fish were tested in pairs with no stimuli. 39 fish were exposed to a single virtual fish while 114 fish were tested in pairs (57 pairs) and without stimuli. When real fish were tested in pairs, data were filtered to only consider cases when the two fish maintained a distance of 0.5 cm to 20 cm between them (tracking accuracy reduced at distances closer than this). Relative 3D positions were then collected by reorganizing the follower's position in the leader's coordinate frame (all relevant filters used in the virtual fish case were also used here). Comparing these two cases—two real fish compared to one real fish swimming with one virtual fish—we find that in the VR, and otherwise, the two fish swim on the same plane (Fig. S17). Hence, all further analyses for decision-making were conducted on this plane in 2 dimensions.

**5.5 Data analysis.** For the symmetric case, the main focus of our analyses was to reveal the role of lateral distance between the virtual fish $L$, on the decision of the real fish to follow these virtual targets. We applied the above mentioned filters to the data and obtained a density plot of the real fish's position in a coordinate system centered at the centroid of the virtual fish's positions (Fig. S15; see Fig. S18 for density plots for varying lateral distance between the virtual fish). The marginal distributions along the direction of the virtual fish's motion for various lateral distances $L$ are then normalized (proportion of maximum) and stacked to obtain the bifurcation plot (Fig. S15). This figure plot shows the effect of virtual fish geometry on the real fish's position while following them. An identical protocol was followed to obtain the three-choice bifurcation plot. For the asymmetric case, we obtain the probability density function of the position of the real fish along the axis perpendicular to its direction of motion. As predicted by our model, the larval zebrafish predominantly assumes one of two positions behind its virtual conspecifics (see main Fig. 4 for comparison of model to data).

### Supplementary Text

#### Experiments in virtual reality

Testing our model predictions experimentally is expected to be difficult. If we are correct, by far the clearest window into the system dynamics will be when animals are presented with two, or more, identical options. This is due to the fact that the very reason that the brain should exhibit bifurcation dynamics—to maximize sensitivity—will also result in amplification of subtle differences between options to obscure our ability to see the underlying system bifurcations. The fact that the experimentalist may often be unaware of such differences (such as a slight air motion, or light gradient, or other differences imperceptible to humans), and that these differences can break the symmetry (between apparently identical options) makes these experiments extremely challenging with a conventional design. To address this limitation, we conduct our experiments in immersive virtual reality (46) in which we can (instantly) randomize our starting conditions, and conduct relatively high-throughput analysis of spatial decision-making.

#### Predictions for symmetric geometries and increasing number of targets

So far, we have discussed predictions from the model and experimental results for the two- and three-choice cases for specific geometries. However, as described section 5.4, we conducted choice experiments for fruit-flies in three different geometrical configurations. An especially interesting case here is one where the targets are in radial symmetry—two targets 180° apart or three targets 120° apart. Once again, we find congruence among predictions of our neural model, the animal collectives model and behavioral experiments with flies (Fig. S9). Because these symmetric conditions represent cases where the animal is already beyond the bifurcation angle, we find that it goes straight to one of the available targets. Further, to illustrate model results beyond three targets, we also ran simulations for four, five, six, and seven targets. Once again, our predictions hold and the agent continues to eliminate targets based on egocentric geometry, thus binarising its decisions (see Fig. 2 in main text).

#### A mental representation of space

Based on direct comparison with experimental results from fruit-flies (*Drosophila melanogaster*), desert locusts (*Schistocerca gregaria*) and zebrafish (*Danio rerio*), we conclude that "cosine-shaped" interactions cannot explain trajectory patterns observed in real animals; that the brain represents space in a non-Euclidean fashion and excitatory interactions among neurons are more local (Fig. S1). Beyond this, the bifurcation patterns observed are agnostic to the exact nature of neural interactions. We illustrate this by using truncated Ricker wavelet (Mexican hat-shaped) neural interactions that produce similar predictions at the level of animal movement (Fig. S1). We specifically chose this function as it has been shown to represent orientation selectivity in neurons of the visual cortex (52).

$$J_{ij} = A\left(1 - h\theta_{ij}^2\right)\exp(-h\theta_{ij}^2) - c \qquad [57]$$

where, $A$ is the amplitude of the Ricker wavelet/Mexican hat, $\theta_i j$ is the angle between preferred directions of spins $i$ and $j$, $h$ is the concentration of the hat and $c$ represents global inhibition. Thus, neural interactions with both long-range inhibition and global inhibition make similar predictions regarding the animal's movement.

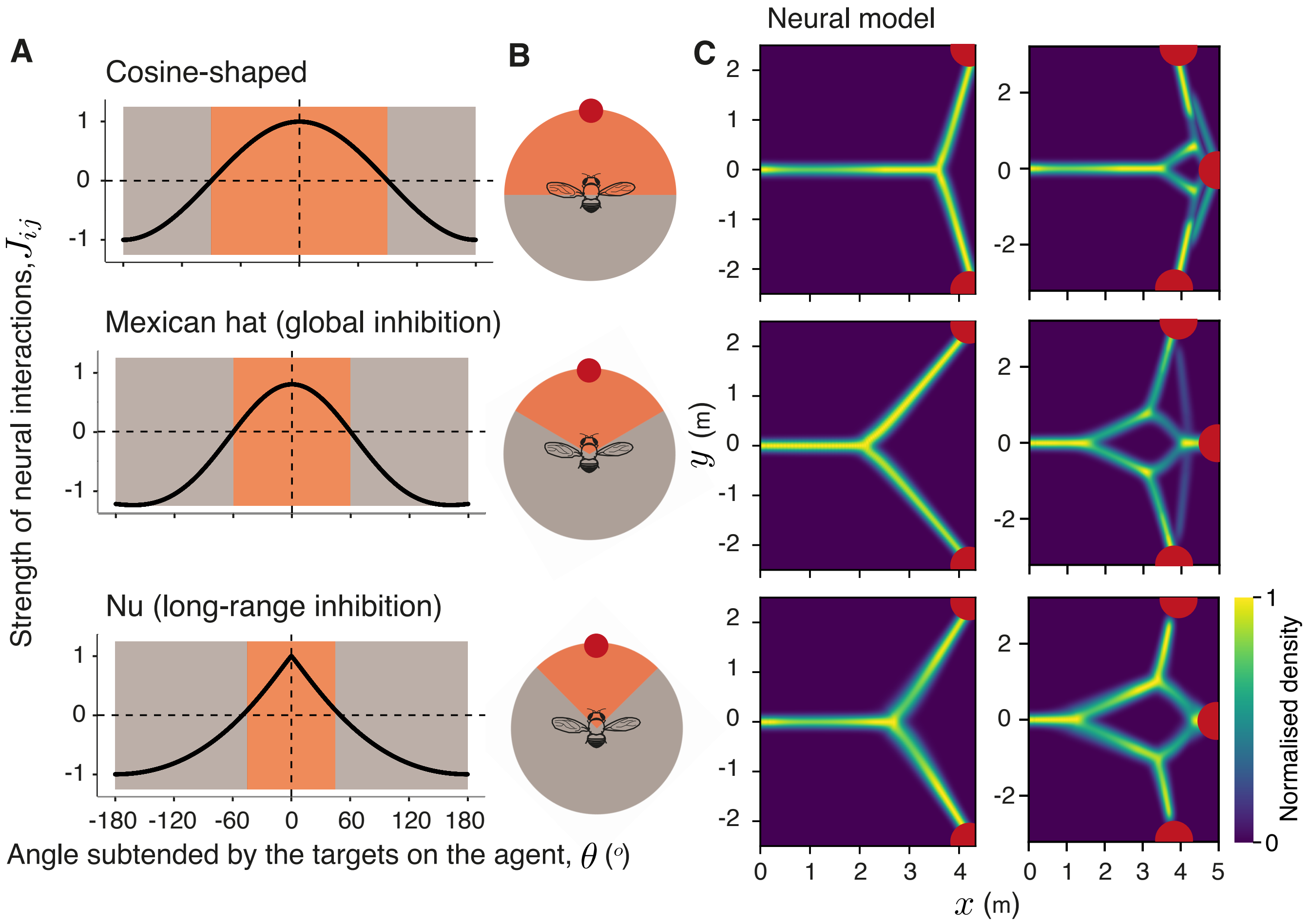


**Fig. S1.** Relationship between the nature of neural interactions and the bifurcation patterns observed in the agent's trajectory. (A) The strength of neural interactions as a function of the angular distance between directions encoded by the spins. We explore models with three different type of neural interactions, a global "cosine-shaped" interactions model, and two local models—with a truncated Mexican hat-shaped interaction curve (amplitude $A = 1.8$, hat concentration $h = 0.25$ and global inhibition $c = 1$), and a model where $\nu = 0.5$. The orange region indicates angles where $J_{ij}$ is positive—excitatory interactions, and the grey region indicates negative $J_{ij}$—inhibitory interactions. (B) A simplified representation of this in polar coordinates. (C) Density plots of trajectories adopted by a model for the corresponding neural interaction curves. The axes represent $x-$ and $y-$coordinates in Euclidean space. Comparing agent trajectories from the simulations with experimental results (see Fig. 1 in main text) from fruit flies and desert locusts reveals a lack of fit by a model with "cosine-shaped" interactions.

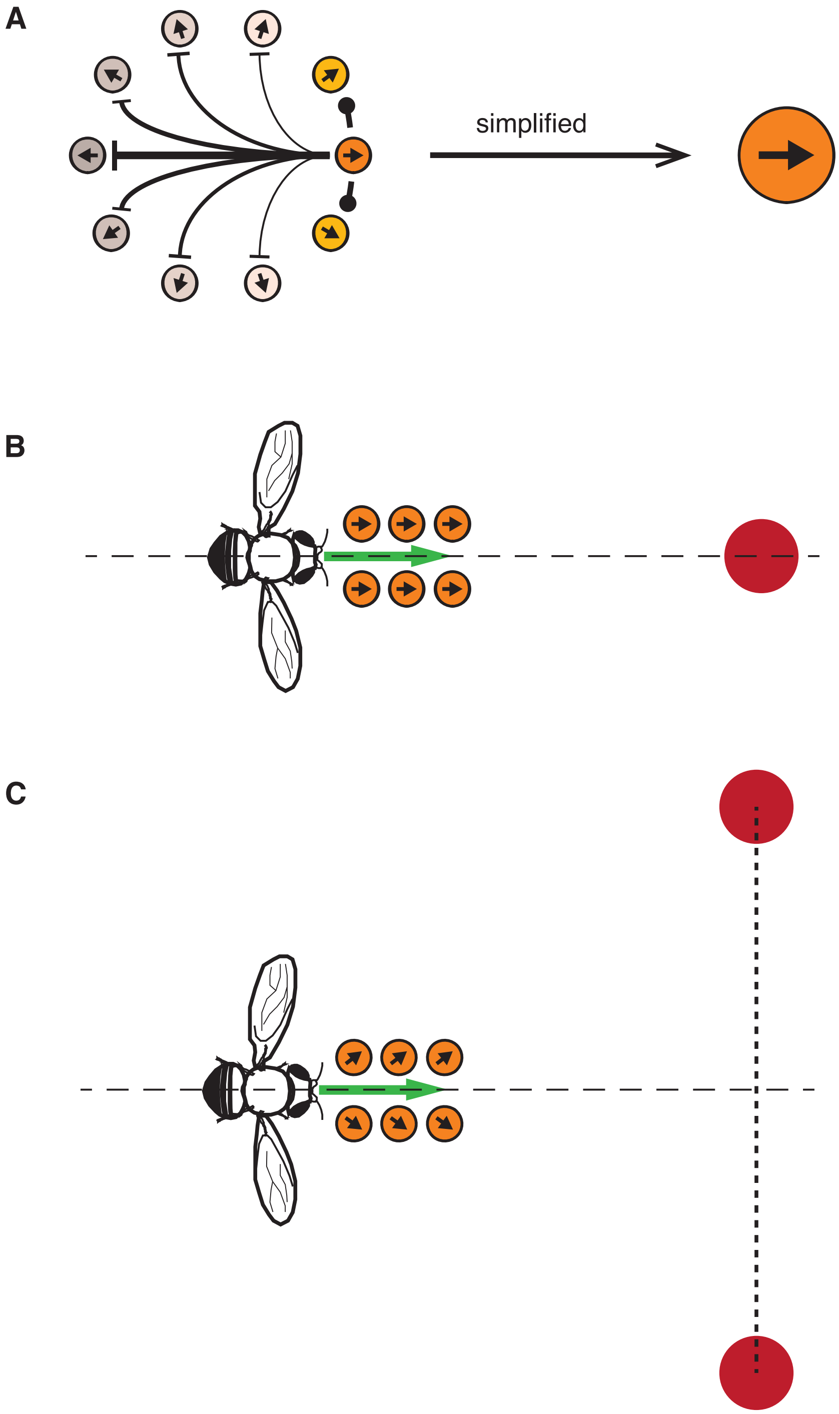


**Fig. S2.** Schematic of the goal-direction cells in an animal's brain when moving through space. (A) It is known that an ensemble of neurons in the brain collectively represents direction to goals (33). Here, we simplify this by representing spins in the model that individually represent direction to a target. (B) The decision-making ensemble is assumed to be a collection of such goal-directed spins. (C) In presence of multiple targets, the ensemble is composed of multiple neural populations that encode directions to the different targets. The agent's decision is thus a consensus among spins in this ensemble.

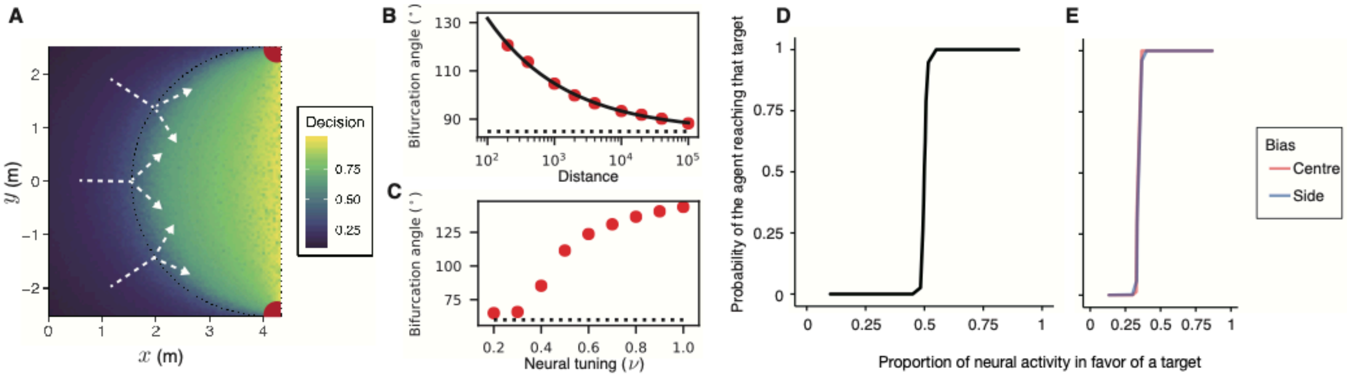


**Fig. S3.** Robustness of bifurcations to spatial and system parameters. (A) A heatmap showing whether or not the neural network has reached a consensus. The agent makes a decision close to the black dotted arc, which represents the locus of a point that is equiangular from the two targets. Angle subtended by the targets on the animal determines the location of the bifurcation. White dotted lines show example trajectories of animals as they would move through space. The axes represent $x-$ and $y-$coordinates in Euclidean space. The exact value of the critical bifurcation angle results from the interplay between two timescales—a timescale for movement and one for the neural dynamics. (B) Effect of the starting distance to the targets on the critical bifurcation angle. We fit an exponential decay to the points to obtain the critical angle (represented here as a dotted line). (C) The neural tuning parameter $(\nu)$ also influences the bifurcation angle. Here, the angle flattens at 60° (represented by the dotted line) as this is the starting angular condition where the agent is initialized. (D-E) Minor difference between the targets causes the agent to choose the correct target with near certainty. The slope of the sigmoid indicates sensitivity of the system. D shows this sensitivity in the presence of two targets while E shows this for the three-target case. Here, we separate sensitivity to the center target vs sensitivity to a side target. As shown, the agent is equally sensitive to all three targets in its environment. See Table S1 for model parameters used here.

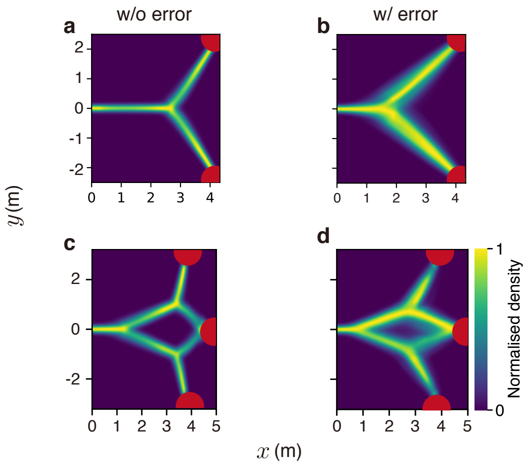


**Fig. S4.** The role of directional error on simulated trajectories. Results of our neural model without (A, C) and with (B, D) implementation of gaussian error and the 'overlap' function for decision-making in a two- (A and B) and three-choice (C and D) context. The symmetric case modelled here puts the targets at an angular configuration where the 'overlap' function does not affect predicted results. However, Gaussian error still introduces noise in the simulated trajectories. Parameters used were identical to Fig. S8. The axes represent $x-$ and $y-$coordinates in Euclidean space.

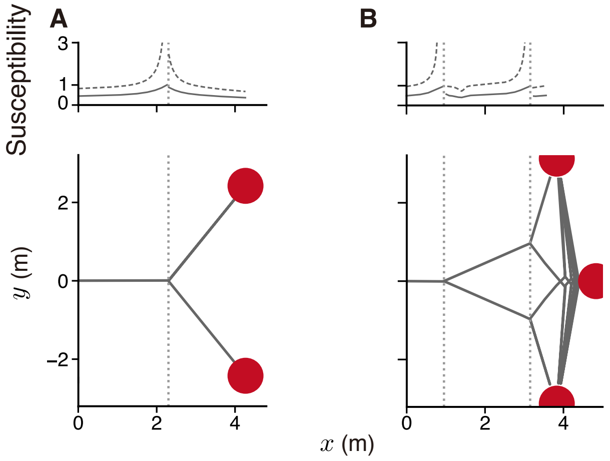


**Fig. S5.** Predicted trajectories and susceptibility from the mean-field approximation. Predicted animal trajectories for decision-making in a two-choice (A) a three-choice (B) context. In both A and B, the susceptibility in the mean-field approximation (dashed line) diverges at the bifurcation points (represented here by the dotted vertical line). The solid line represents the short time average responses of one spin flip in reaction to one extra spin towards one of the targets. The short time response shows a peak at the bifurcation point and is normalized by its maximal value. See Table S1 for parameters used.

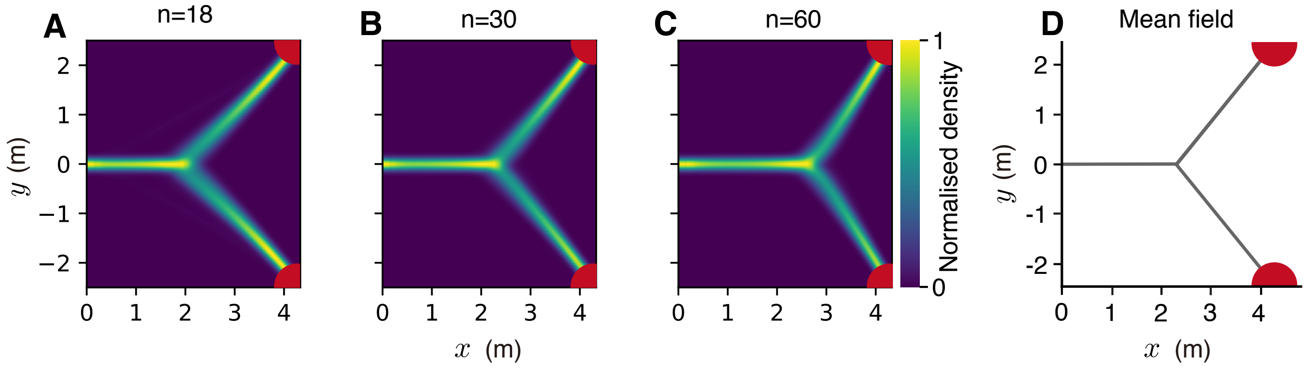


**Fig. S6.** Effect of system size on observed bifurcations. Comparison of panels A-D shows that bifurcation patterns are observed both for small and large system sizes. Panels A, B, and C show trajectories predicted by the neural model for system sizes $N = 18$, $N = 30$, and $N = 60$ respectively. Panel D shows the predicted trajectories at the mean-field limit of very large system sizes $N \to \infty$. The axes represent $x-$ and $y-$coordinates in Euclidean space.

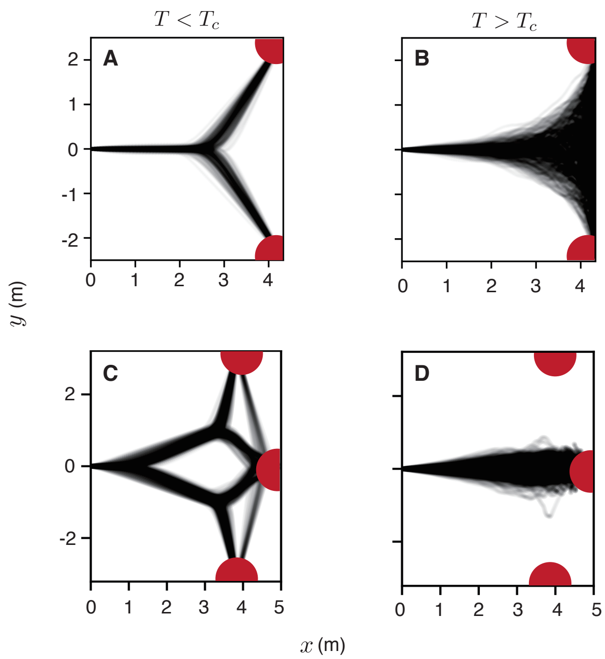


**Fig. S7.** Role of neural noise on producing the experimentally observed bifurcation patterns. Each panel contains trajectories from 500 replicate simulations of our neural decision-making model. The model reproduces experimentally observed bifurcation patterns below a critical level of neural noise (A and C). Above this critical noise level, the agent exhibits diffusive movement for the two-choice case (B) with a bias towards more central targets (D). A and B show trajectories for decision-making in the presence of two targets, while C and D show trajectories for decision-making in the presence of three targets. The axes represent $x-$ and $y-$coordinates in Euclidean space. See Table S1 for parameter values used in A and C. B and D were produced with identical parameters except the neural noise parameter $T = 2.0$.

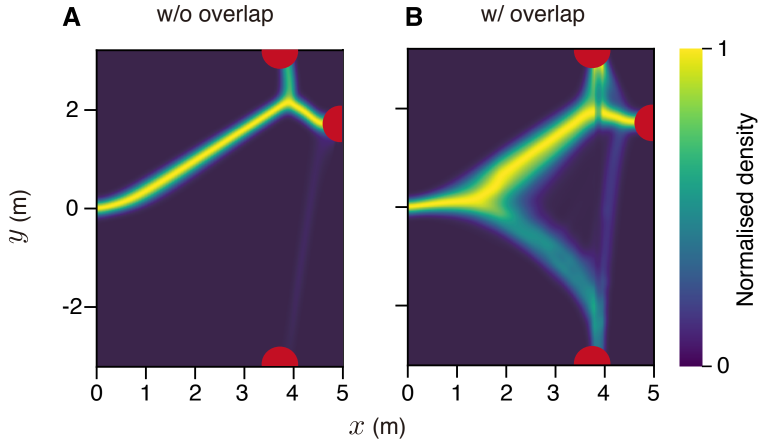


**Fig. S8.** Decision-making in asymmetric geometries. Results of our neural model without (A) and with (B) implementation of the 'overlap' function. This function allows us to account for spatial geometries where some targets may be in directional proximity compared to others. The Gaussian error in B was of standard deviation $\sigma_\theta = 0.25$. The axes represent $x-$ and $y-$coordinates in Euclidean space.

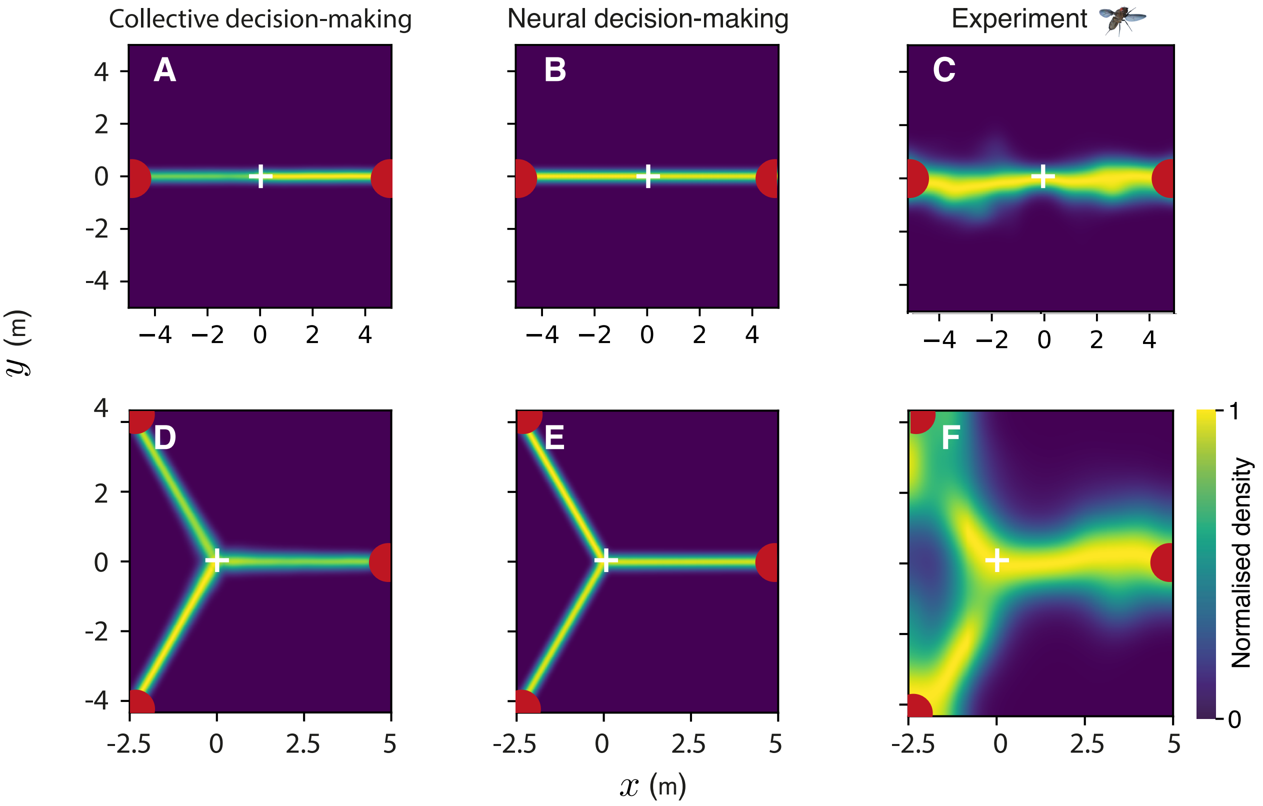


**Fig. S9.** Decision-making in radial symmetry. Trajectories predicted by two different models of decision-making and experimental trajectories obtained from fruit flies, all in the presence of two and three targets placed in radial symmetry. The animal starts at (0, 0) and chooses one of the available targets. The axes represent $x-$ and $y-$coordinates in Euclidean space. Panels A and D show results for two- and three-choice decision making from a collective decision-making model, B and E show results from our neural decision-making model, and C and F show experimental results of fruit-flies exposed to two and three identical targets respectively. See Table S1 for parameters used in A and D, and Table S2 for parameters used in B and E.

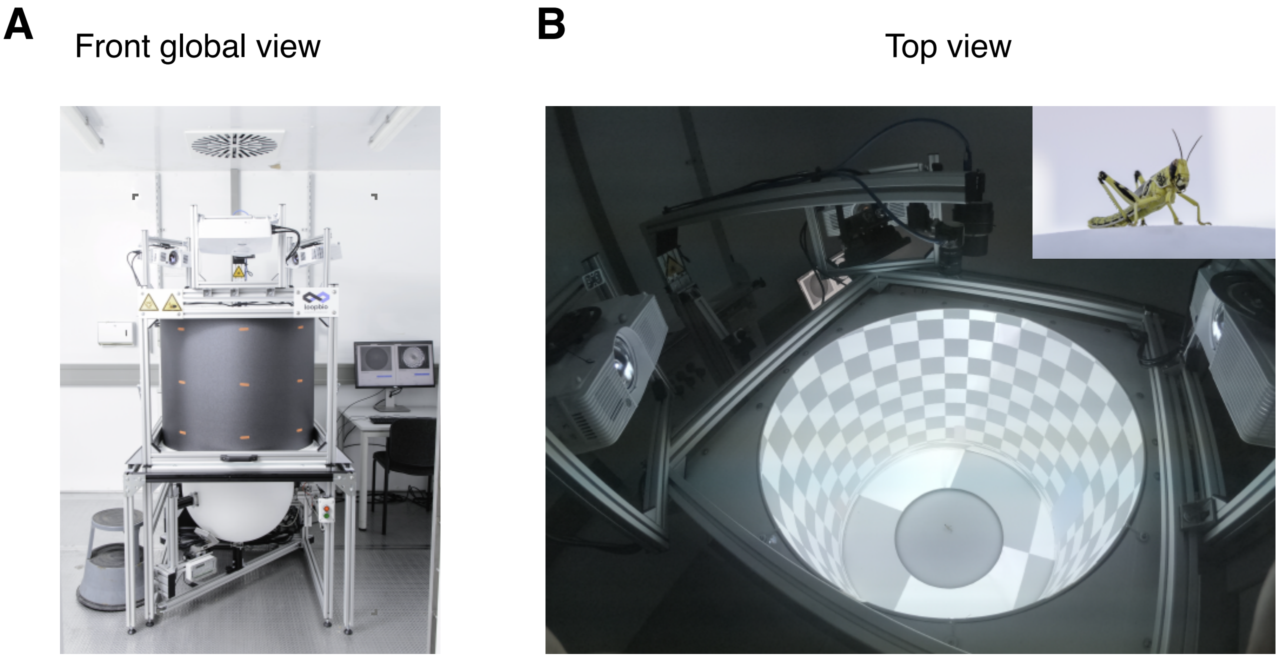


**Fig. S10.** Overview of the locustVR experimental setup. (A) The front-view of the locustVR setup shows the three projectors, the cylindrical projection surface, and the locomotion compensator sphere. (B) The top-view of the setup shows a custom checkerboard stimulus along with the locust on the center of the locomotion compensator sphere. The inset shows a zoomed in side-view of the locust.

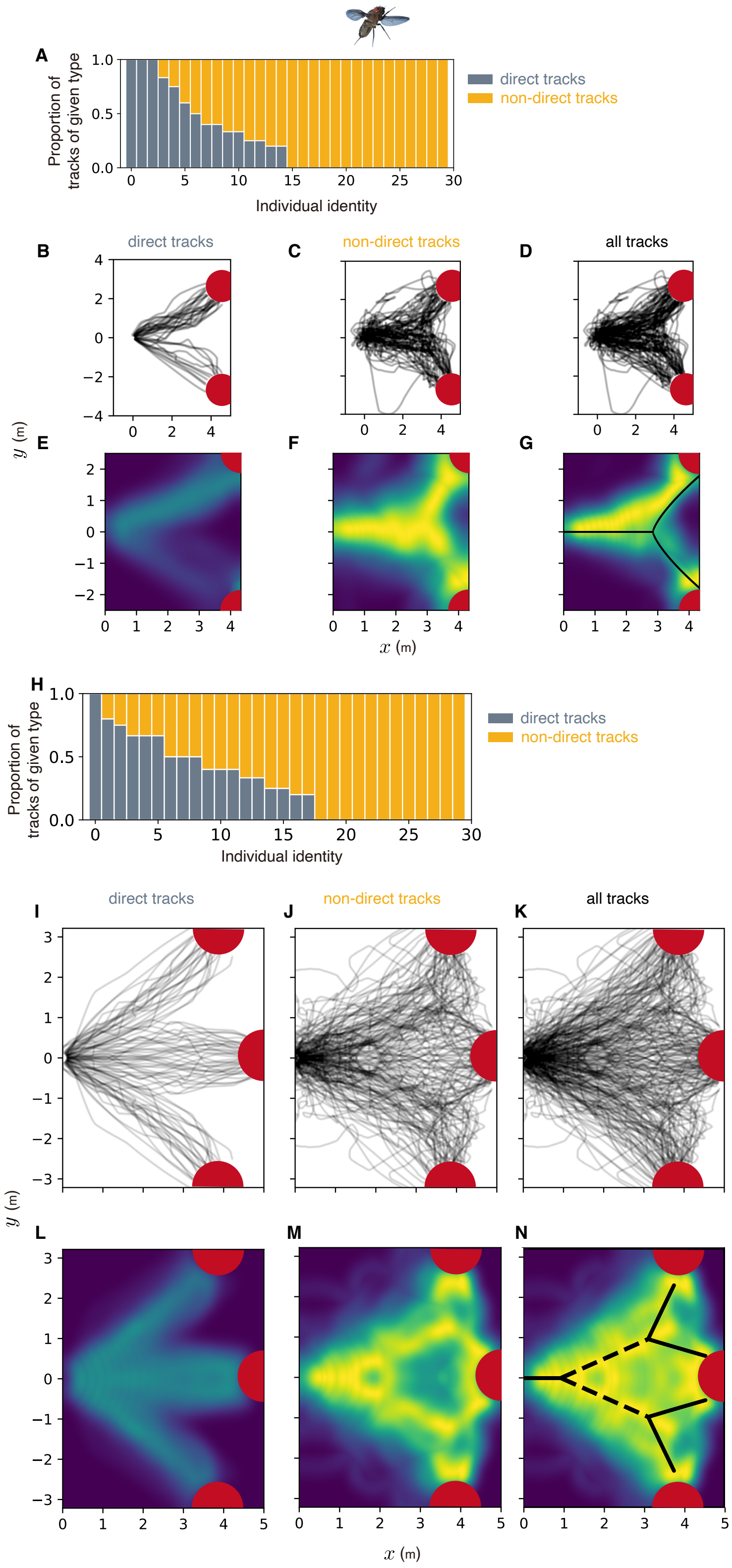


**Fig. S11.** Characterisation of fruit fly tracks at the individual and population level when exposed to two and three targets. A and H show consistency in individual behavioral tendencies in exhibiting either direct or non-direct tracks. (B-G) present raw trajectories (B-D) and density plots (E-G) of fruit flies exposed to two targets. The axes represent $x-$ and $y-$coordinates in Euclidean space. A shows trajectories where the fruit fly reached the target in a relatively short duration, where trajectories to the targets were relatively direct, and B shows the remaining trajectories. E and F show the corresponding density plots, normalized such that the maximum intensity in F is set to 1, and in E is set to the proportion of trajectories in F relative to this condition. D and G show the raw trajectories and the normalized density plot for all fly experiments combined. Similarly, I-N present raw trajectories (I-K) and density plots (L-N) of fruit flies exposed to three targets. I and L show direct trajectories to a target and the corresponding density plot. J and M show the remaining trajectories that potentially exhibit the bifurcations, and the corresponding density plot. K and N show raw trajectories and the density plot for all fly experiments combined. Note that the three-choice trajectories here are symmetrized for the sake of visualization. Our conclusions do not differ when we include all the fly data. To show this, we fit the piecewise phase transition function (shown in black) to density plots G and N.

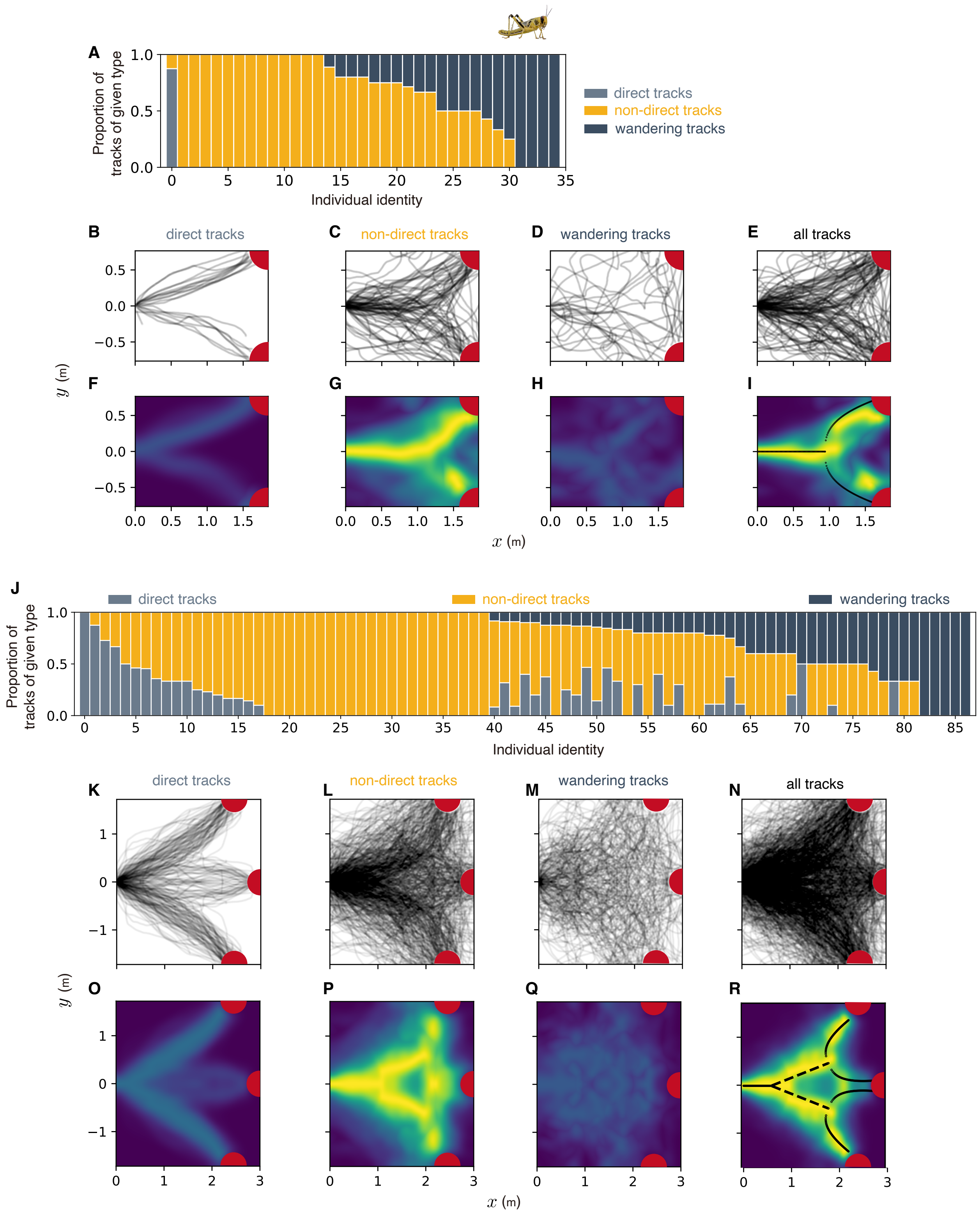


**Fig. S12.** Characterisation of desert locust tracks at the individual and population level when exposed to two and three targets. A and J show consistency in individual behavioral tendencies in exhibiting either direct, non-direct or wandering tracks. (B-I) present raw trajectories (B-E) and density plots (F-I) of locusts exposed to two targets. The axes represent $x-$ and $y-$coordinates in Euclidean space. B shows trajectories where the locust reached the target in a relatively short duration, where trajectories to the targets were relatively direct, D shows trajectories where the locust took long to reach the target, where trajectories to the targets were noisy, and C shows the remaining trajectories. F-I show the corresponding density plots, normalized such that the maximum intensity in G is set to 1, and in F and H is set to the proportion of trajectories in G relative to this condition. E and I show the raw trajectories and the normalized density plot for all locust experiments combined. Similarly, K-R present raw trajectories (K-N) and density plots (O-R) of locusts exposed to three targets. K and O show direct trajectories to a target and the corresponding density plot, M and Q show noisy trajectories to a target and the corresponding density plot, and L and P show the remaining trajectories that potentially exhibit the bifurcations, and the corresponding density plot. N and R show raw trajectories and the density plot for all locust experiments combined. Note that the three-choice trajectories here are symmetrized for the sake of visualization. Our conclusions do not differ when we include all the locust data. To show this, we fit the piecewise phase transition function (shown in black) to density plots I and R.

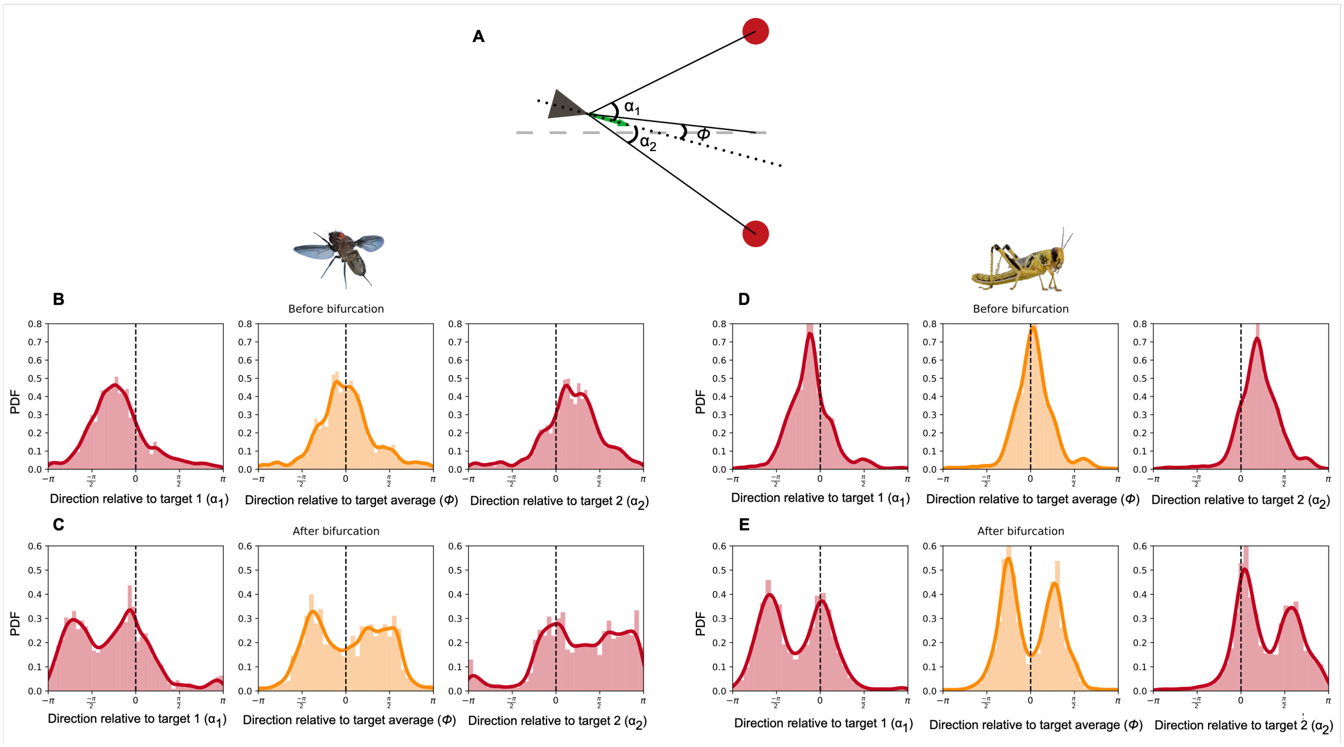


**Fig. S13.** Characterization of the direction of movement of fruit-flies and desert locusts in the non-direct tracks. (A) A schematic representation of the various reference frames used to characterize the animals' direction of movement—the green arrow represents the animals' velocity vector, the black dotted line is their current direction of movement, $\alpha_1$ and $\alpha_2$ represent movement direction relative to the two targets and $\phi$ represents movement direction with respect to the average of the egocentric target directions. The grey dashed line is the perpendicular bisector of the line connecting the two targets. All animals start their experimental trial at a fixed distance from the targets on this line. (B and D) Before the bifurcation, both flies and locusts tend to direct their movement directly towards the average of the egocentric target directions ($\phi$). Thus, they do not fixate on either target. Beyond the critical angular difference, however, (C and E) they switch from directing their motion towards the average of the target directions, to fixating on one, or other, of them and thus exhibiting directed motion towards the selected target (in the direction of $\alpha_1$ or $\alpha_2$ to target 1 or target 2 respectively). The second peak evident when this occurs, offset from the heading by a large angle, shows the direction towards the unselected target.

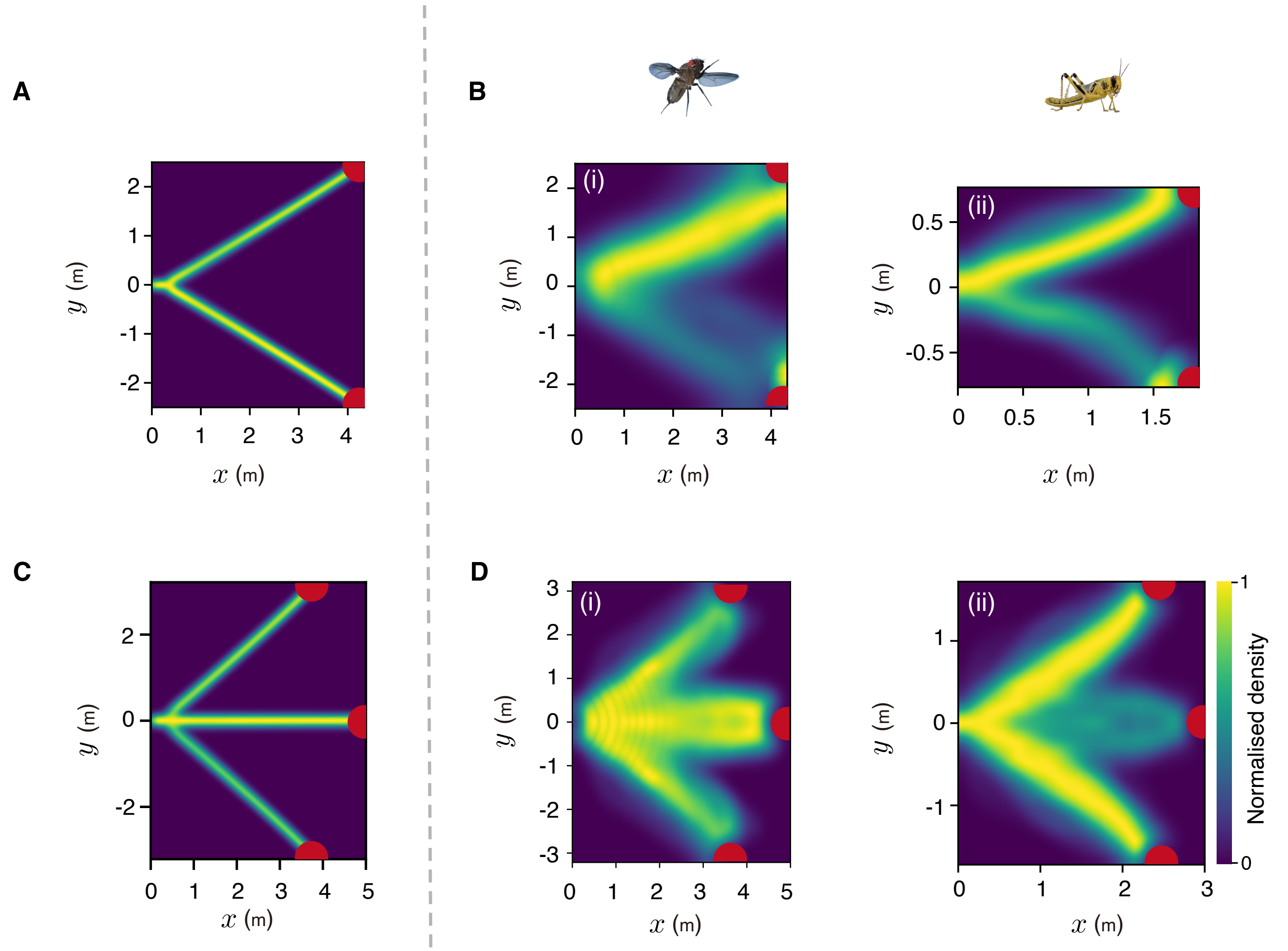


**Fig. S14.** Model results where the animal is predicted to directly approach one of the presented targets and corresponding trajectories from experiments with fruit flies and desert locusts. (A and C) Density plots of predicted trajectories for two and three-choice decision-making with high neural tuning ($\nu = 0.2$). Under these parameter conditions, the model predicts that the animal should directly move towards one of the presented targets. (B and D) show density plots of corresponding direct tracks exhibited by fruit flies and desert locusts respectively.

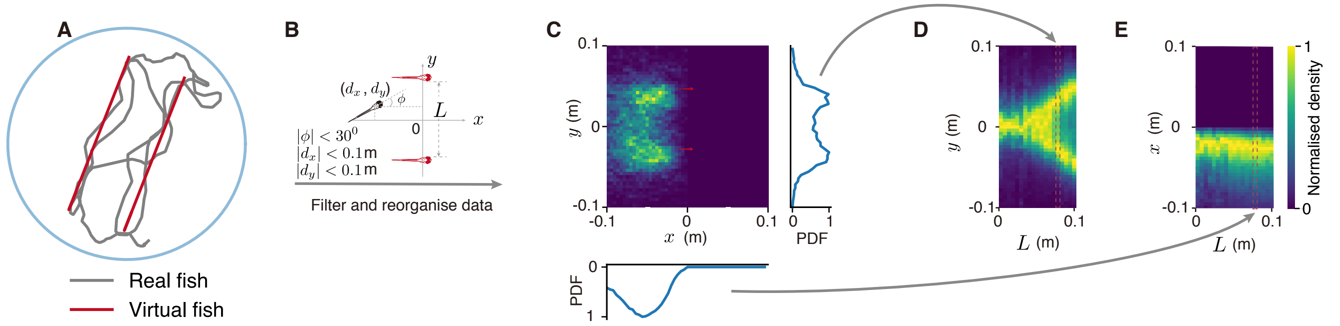


**Fig. S15.** Overview of experiments and data analyses with larval zebrafish. (A) Two virtual fish swim back-and-forth in the arena beside each other at varying lateral distances (0.08 m in this particular case). (B) Data were centered on a coordinate system with origin at the centroid of the virtual fish's positions and decisions were considered along the axis perpendicular to their direction of motion. (C) An example of the real fish's position density relative to the virtual fish. We obtain a normalized marginal probability distribution of the real fish's position (perpendicular to the virtual fish's movement direction) and stack these distributions for varying lateral distances between the virtual fish (D). We can do this without losing information along the virtual fish's movement direction as the real fish maintains relatively stable front-back distance with its virtual conspecifics (E).

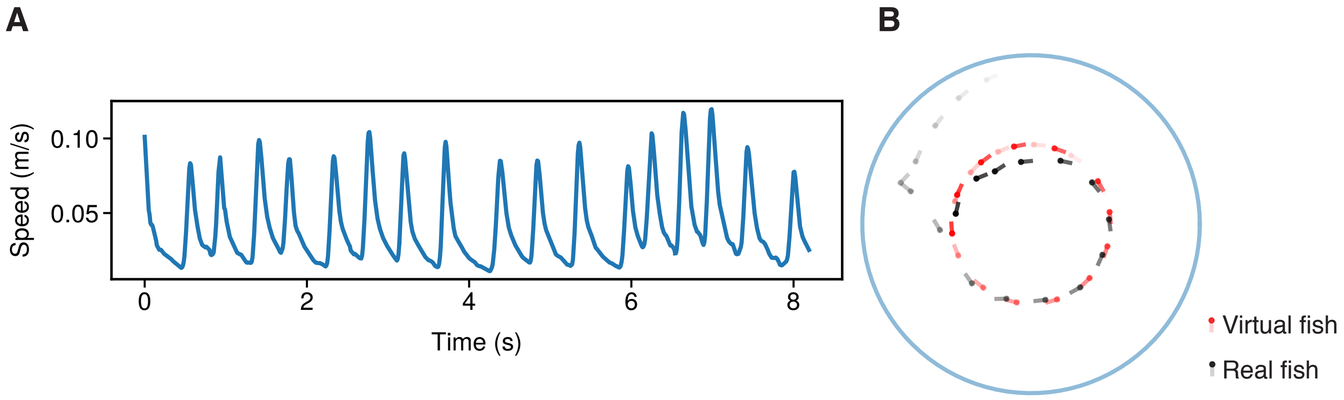


**Fig. S16.** Behavioral experiments with larval zebrafish in virtual reality. (A) Burst-and-glide movement adopted by the virtual zebrafish. The average movement speed was extracted statistically from freely swimming larval zebrafish and the burst-and-glide movement was adopted from a random real fish. (B) Interaction of the real fish with its virtual conspecific in one of the control runs where the virtual fish swims in a circular path.

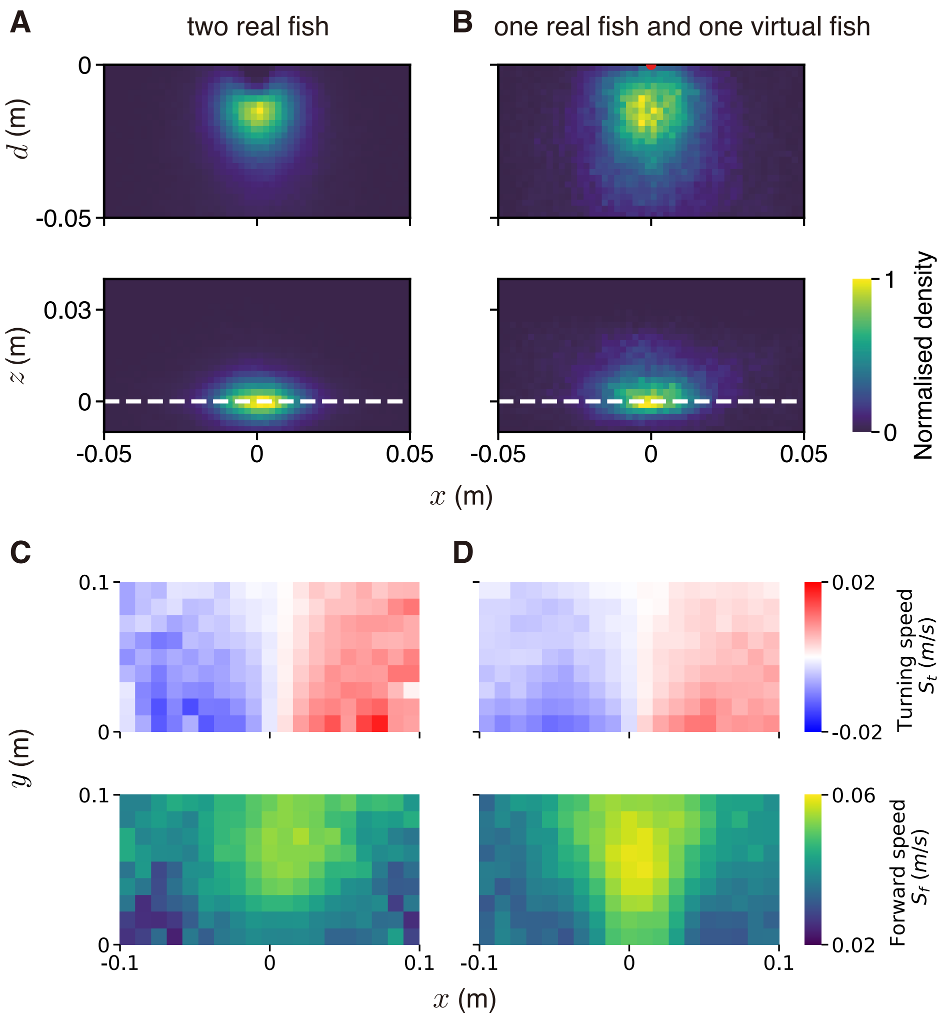


**Fig. S17.** Fish respond similarly to real and virtual conspecifics. Density plots from 3D tracking of pairs of fish within and without the VR. Both panels A and B represent the position of a follower in a coordinate system centered on the leader. The dashed line in B represents position of the leader on the $z-$plane. Panels C and D show the turning (left-right) and the forward (front-back) speeds of the follower in a coordinate system centered on the leader. (A and C) Two larval zebrafish fish swimming together. (B and D) A larval zebrafish swimming with a virtual conspecific.

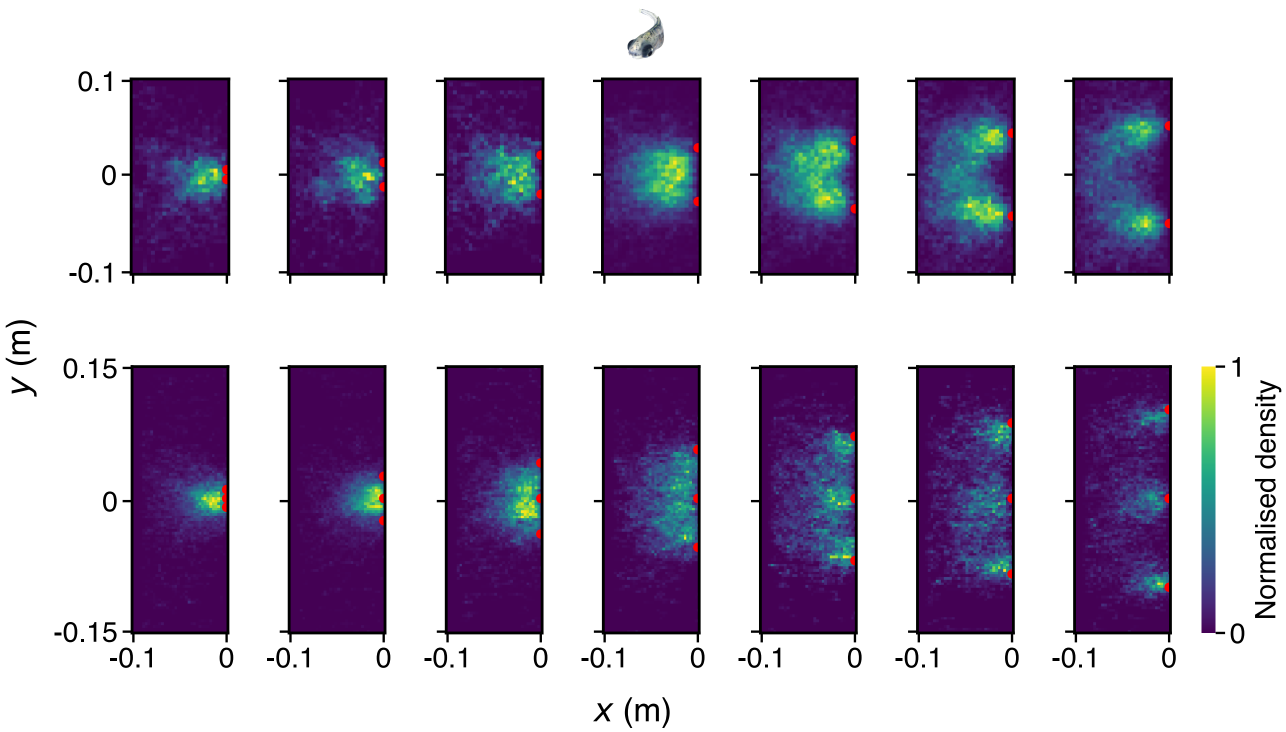


**Fig. S18.** Normalized density plots for larval zebrafish exposed to two or three virtual fish. The red dots represent positions of the virtual zebrafish that are presented to the real fish. Density plots represent position of the real fish as it follows these virtual fish. Top row presents positions of the real fish in presence of two virtual fish and the bottom row presents positions of the real fish in presence of three virtual fish.

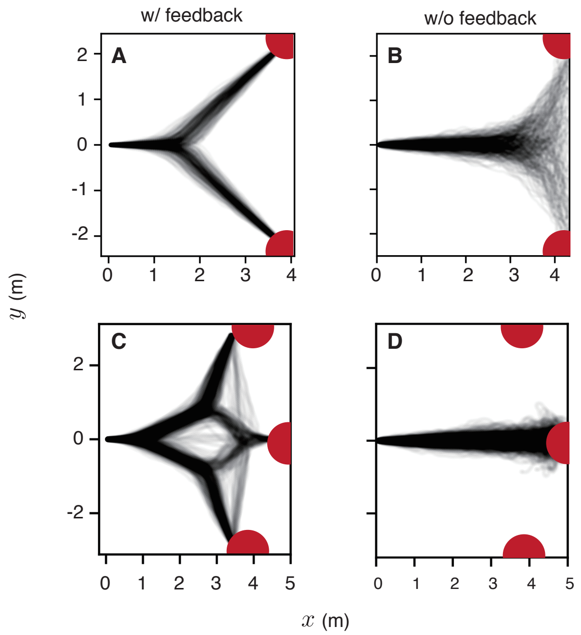


**Fig. S19.** Role of feedback in producing the experimentally observed bifurcations in a model of animal collectives. Trajectories resulting from a decision-making model of animal collectives with (A and C) and without feedback (B and D) on individual preferences (500 replicates). A and B show trajectories for decision-making in the presence of two targets, while C and D show trajectories for decision-making in the presence of three targets. The axes represent $x-$ and $y-$coordinates in Euclidean space. See Table S2 for parameters used in A and C. For B and D, we removed explicit feedback on individual preferences by setting $\omega_{inc} = 0$ and $\omega_{dec} = 0$. However, with no feedback, the group will almost always split into subgroups that all go to their preferred target. Hence, B and D also included uninformed individuals that function to keep the group together. In these simulations, each target was preferred by 12 individuals and the rest were uninformed—simulations with two options had 36 uninformed individuals out of 60 while the simulations with three options had 24 uninformed individuals.

**Table S1. An overview of the parameter values explored for the neural decision-making model.**

| Parameter | Symbol | Value(s) |
|---|---|---|
| System size (simulations) | $N$ | 60 |
| Neural tuning | $\nu$ | 0.5 |
| Directional noise (simulations) | $\sigma_e$ | 0.02 |
| Neural noise / Temperature | $T$ | 0.2 |
| Lateral distance in cm (fish simulations) | $L$ | 0–10 |
| Front-back distance in cm (fish simulations) | $d$ | 1.5–3 |
| Directional error (overlap implementation) | $\sigma_\theta$ | 0.25 |

**Table S2. An overview of the parameter values explored for the collective decision-making model.**

| Parameter | Symbol | Value(s) |
|---|---|---|
| Total agents | $N$ | 60 |
| Repulsion radius | $r_r$ | 1 |
| Attraction / Alignment radius | $r_a$ | 6 |
| Turning rate | $\psi$ | 2 |
| Speed | $\lvert v\rvert$ | 1 |
| Omega initialization | $\omega_0$ | 0.3 |
| Omega increment | $\omega_{inc}$ | 0.012 |
| Omega decrement | $\omega_{dec}$ | 0.0008 |
| Maximum omega | $\omega_{max}$ | 0.4 |
| Timestep increment | $\Delta t$ | 0.1 |
